# A ciliopathy complex builds distal appendages to initiate ciliogenesis

**DOI:** 10.1101/2021.03.01.433418

**Authors:** Dhivya Kumar, Addison Rains, Vicente Herranz-Pérez, Quanlong Lu, Xiaoyu Shi, Danielle L. Swaney, Erica Stevenson, Nevan J. Krogan, Bo Huang, Christopher Westlake, Jose Manuel Garcia-Verdugo, Bradley Yoder, Jeremy F. Reiter

## Abstract

Cells inherit two centrioles, the older of which is uniquely capable of generating a cilium. Using proteomics and super-resolved imaging, we identified a module which we term DISCO (DIStal centriole COmplex). DISCO components CEP90, MNR and OFD1 underlie human ciliopathies. This complex localized to both distal centrioles and centriolar satellites, proteinaceous granules surrounding centrioles. Cells and mice lacking CEP90 or MNR did not generate cilia, failed to assemble distal appendages, and did not transduce Hedgehog signals. Disrupting the satellite pools did not affect distal appendage assembly, indicating that it is the centriolar populations of MNR and CEP90 that are critical for ciliogenesis. CEP90 recruited the most proximal known distal appendage component, CEP83, to root distal appendages formation, an early step in ciliogenesis. In addition, MNR, but not CEP90, restricted centriolar length by recruiting OFD1. We conclude that DISCO acts at the distal centriole to support ciliogenesis by restraining centriole length and assembling distal appendages, defects in which cause human ciliopathies.

**eTOC summary:** Kumar et al. identifies a multi-protein complex called DISCO (DIStal centriole COmplex) required to nucleate distal appendages and restrain centriole elongation, essential for the initiation of cilium assembly. Without DISCO, cells fail to ciliate and transduce Hedgehog signals, critical for mammalian development.

## INTRODUCTION

Centrioles are ancient, microtubule-based structures with two main functions: first, they are core components of the centrosome, the primary microtubule organizing center, and, second, they are the foundations for cilia, cellular antennae specialized for signaling. While the main barrel of each centriole is comprised of nine radially arranged microtubule triplets, many proteins apart from tubulin comprise the centrioles (Winey and O’Toole, 2014; Keller et al., 2005, 2009; Jakobsen et al., 2011; Andersen et al., 2003). To date, the parts list of centrioles is incomplete.

Centriole structure can vary between species, but within a particular cell type, centriole length is nearly uniform, as are centriolar number and timing of assembly (Goehring and Hyman, 2012; Kong et al., 2020). Coordination with the cell cycle entrains this uniformity: centrioles duplicate during interphase and, at mitosis, each daughter cell inherits an older, mother centriole and a younger, daughter centriole (Nigg and Holland, 2018; Breslow and Holland, 2019).

The differences in centriolar age dictates different structures and functions. Only the mother centriole can assemble a cilium, and only it possesses subdistal and distal appendages. Through the distal appendages, the mother centriole attaches to pre-ciliary vesicles, an early step in ciliogenesis (Tanos et al., 2013; Sillibourne et al., 2013; Schmidt et al., 2012b). In contrast, sub-distal appendages are dispensable for ciliogenesis (Tanos et al., 2013; Mazo et al., 2016; Chong et al., 2020). Although components of appendages have been identified (Bowler et al., 2019; Yang et al., 2018; Chong et al., 2020), how they are assembled remains enigmatic.

Related to their roles in fundamental cellular processes such as cell division and intercellular communication, centriole dysfunction causes diverse human developmental disorders including ciliopathies and microcephaly (Nigg and Holland, 2018; Reiter and Leroux, 2017). For example, mutations in *CEP90* (*Centrosomal protein of 90 kDa*, also known as *PIBF1*), *MNR* (*MOONRAKER*, also known as *KIAA0753* or *OFIP*) or *OFD1* cause Joubert syndrome (JBTS), a ciliopathy characterized by brainstem and cerebellar malformations (Wheway et al., 2015; Shen et al., 2020; Hebbar et al., 2018; Stephen et al., 2017; Hammarsjö et al., 2017; Coene et al., 2009). Similarly, in many cancers, the number or structure of centrioles is dysregulated (Gönczy, 2015; Marteil et al., 2018).

CEP90 has been identified as a component of centriolar satellites, protein assemblies that orbit the centrosome (Kim et al., 2012; Kodani et al., 2015). Knockdown of *CEP90* alters the localization of centriolar satellites, which has been proposed to compromise ciliogenesis (Kim and Rhee, 2011; Kim et al., 2012).

We have used a combination of label retention expansion microscopy and structured illumination microscopy (LR-ExSIM) (Shi et al., 2019) and found that, in addition to being a component of centriolar satellites, CEP90 is a component of the distal end of centrioles. Proteomic analyses revealed that CEP90 is part of an evolutionarily conserved complex, which we term DISCO (DIStal centriole COmplex) that includes MNR and OFD1. Mouse genetic studies revealed that CEP90 and MNR are key regulators of mother centriole function, essential for normal vertebrate development, Hedgehog signaling and ciliogenesis. In investigating how CEP90 and MNR function in ciliogenesis, we found that both are required for the assembly of distal appendages. Mutant analysis demonstrated that the components of this distal centriole complex are recruited to the centriole in a hierarchical manner, culminating in CEP90, which recruits the most proximal distal appendage protein, CEP83, to initiate distal appendage formation. Thus, our work identifies an evolutionarily conserved complex that functions at the distal mother centriole to build distal appendages, an early and critical step in ciliogenesis.

## RESULTS

### CEP90 localizes to centriolar satellites and a ring at the distal ends of centrioles

CEP90 has previously been identified as a component of centriolar satellites (Kim and Rhee, 2011). We confirmed that CEP90 localizes to centriolar satellites by immunostaining human retinal pigment epithelial (RPE1) cells with antibodies to CEP90 and PCM1, a marker of centriolar satellites (Fig. 1a). In addition, a centrosomal pool of CEP90 did not co-localize with PCM1 (Fig. 1a). To more precisely identify where centrosomal CEP90 localized, we dispersed centriolar satellites with nocodazole and found that CEP90 localized with γ-tubulin at centrioles (Fig. 1b). Three-dimensional structured illumination microscopy (3D-SIM) revealed that CEP90 forms rings at centrioles (Fig. 1b).

**Figure 1.**
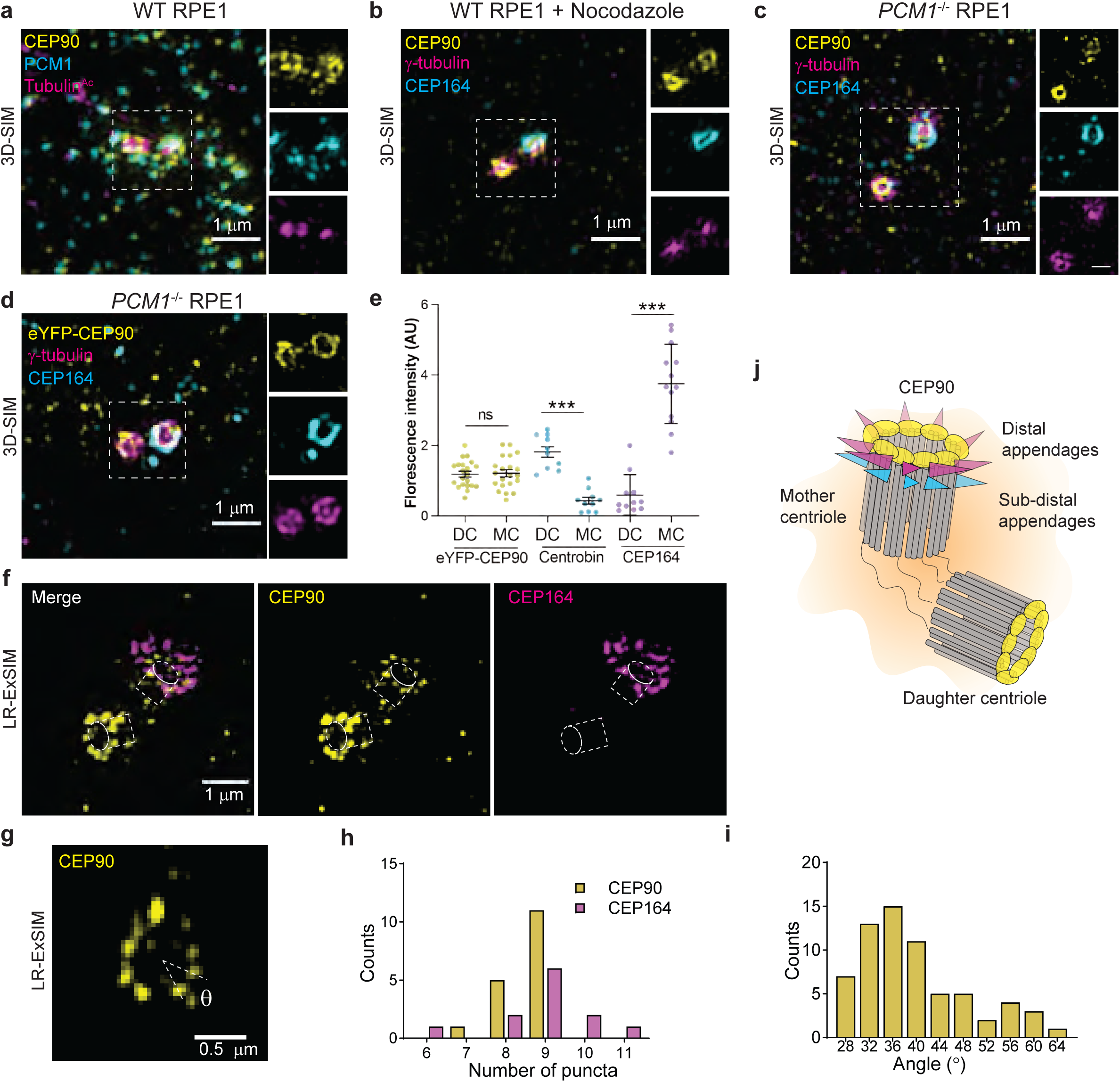
CEP90 localizes to centriolar satellites and the distal centriole. **a.** 3D-SIM of immunostained RPE1 cells reveals localization of CEP90 (yellow) at centriolar satellites (PCM1, cyan) and centrioles (Tubulin^Ac^, magenta) in wild type, cycling RPE1 cells. Scale bar = 1 μm. **b**. Treatment of serum-starved RPE1 cells with nocodazole disperses the centriolar satellites, highlighting 3D-SIM of CEP90 (yellow) rings at centrioles (γ-tubulin, magenta). Distal appendage component CEP164 (cyan) indicates the mother centriole in C and D. Scale bar = 1 μm. **c**. 3D-SIM of serum-starved *PCM1*^-/-^ RPE1 cells shows that CEP90 (yellow) localizes to centrioles (γ-tubulin, magenta) independent of centriolar satellites. Scale bar of main panel and insets = 1 μm and 0.5 μm respectively. **d**. 3D-SIM confirms localization of CEP90 (yellow) at centrioles (γ-tubulin, magenta) in nocodazole-treated, serum-starved eYFP-CEP90 expressing *PCM1*^-/-^ RPE1 cells. Scale bar = 1 μm. **e**. Quantification of eYFP-CEP90, Centrobin and CEP164 fluorescence intensity at daughter (DC) and mother (MC) centrioles from n = 10-20 cells. Horizontal lines indicate means ± SEM. Asterisks indicate p<0.05 determined using unpaired t test and ns = not significant. **f**. LR-ExSIM of RPE1 cells immunostained for CEP90 (yellow) and CEP164 (magenta) reveals that rings of CEP90 are comprised of discrete puncta. CEP90 rings are smaller and more proximal to CEP164 rings. Scale bar = 1 μm. **g**. Example of an LR-ExSIM image of a radially oriented centriole used to quantify the number and angle between adjacent puncta of CEP90. Scale bar = 0.5 μm. **h**. Histogram of number of discrete puncta of CEP90 and CEP164 observed per centriole in LR-ExSIM images. n = 12-17 measurements. **i**. Histogram of the angular spacing between adjoining centriolar CEP90 puncta observed by LR-ExSIM. n = 66 measurements. **j**. Schematic of the ring of CEP90 punctae (yellow), distal appendages (magenta), and sub-distal appendages (blue) at the distal centriole. CEP90 decorates the distal end of mother and daughter centrioles.

Independent of nocodazole, we disrupted centriolar satellites in RPE1 cells by deleting PCM1, a scaffold for centriolar satellites, using CRISPR/Cas9 (Supplementary Fig 1). As described previously, centriolar satellites were not detected in *PCM1*^-/-^ RPE1 cells (Odabasi et al., 2019). Consistent with the loss of satellites, localization of CEP90 to puncta around the centrosome was absent in *PCM1*^-/-^ RPE1 cells (Fig. 1c). As in nocodazole-treated cells, the rings of CEP90 at centrioles remained in *PCM1*^-/-^ RPE1 cells (Fig. 1c), indicating that CEP90 localizes to centrioles independently of PCM1 and centriolar satellites. To confirm antibody staining specificity, we immunostained exogenously expressed eYFP-tagged CEP90 in nocodazole-treated *PCM1*^-/-^ RPE1 cells (Fig. 1d). Localization of eYFP-CEP90 resembled endogenous CEP90 staining, and suggested that CEP90, unlike Centrobin and CEP164, localized to both mother and daughter centrioles with similar fluorescence intensity (Fig. 1e).

Expansion microscopy involves physically expanding samples embedded in a hydrogel (Wassie et al., 2019). To precisely map the localization of CEP90 at centrioles, we combined expansion microscopy together with multi-color label retention and SIM (LR-ExSIM) to minimize signal loss during expansion and provide ∼30 nm lateral resolution (Shi et al., 2019). The distal appendage component CEP164 imaged by LR-ExSIM was comparable to data previously obtained using STORM: CEP164 formed a discontinuous ring at the mother centriole (Shi et al., 2019, 2017). Intriguingly, LR-ExSIM resolved that the ring of CEP90 is comprised of discrete puncta with a mode of nine and separated by a mean angle of ∼ 36 nm (Fig. 1f-i). In agreement with 3D-SIM microscopy, the CEP90 ring at the mother centriole as observed by LR-ExSIM had a smaller diameter (228 ± 23 nm) and was proximal to the distal appendages. Thus, CEP90 comprises a nine-fold ring at the distal centriole (Fig 1j).

### CEP90, OFD1 and MNR form a distal centriole complex

To gain insight into the function of CEP90 at the distal centriole, we proteomically identified CEP90 interactors. More specifically, we generated RPE1 cell lines stably expressing GFP (control) or YFP-CEP90. To discriminate proteins interacting with CEP90 at centriolar satellites and those interacting with CEP90 at centrioles, we also generated *PCM1*^-/-^ RPE1 cell lines stably expressing GFP or YFP-CEP90 in which centriolar satellites are disrupted (Fig. 2a). We immunoprecipitated GFP from these cell lines and detected co-immunoprecipitating proteins by LC/MS/MS. Significance analysis of interactome (SAINTexpress (Teo et al., 2014)) analysis with Bayesian false discovery rate (BFDR) < 0.05 indicated high confidence interactors (Fig. 2b and c). Some CEP90 interactors, such as PCM1, CEP131 and BBS4, were identified in wild type cells, but not in *PCM1*^-/-^ cells, suggesting that these interactions are dependent on centriolar satellites. Consistent with this conclusion, many of these PCM1-dependent interactors, including CEP131 and BBS4, are components of centriolar satellites (Hori and Toda, 2017).

**Figure 2.**
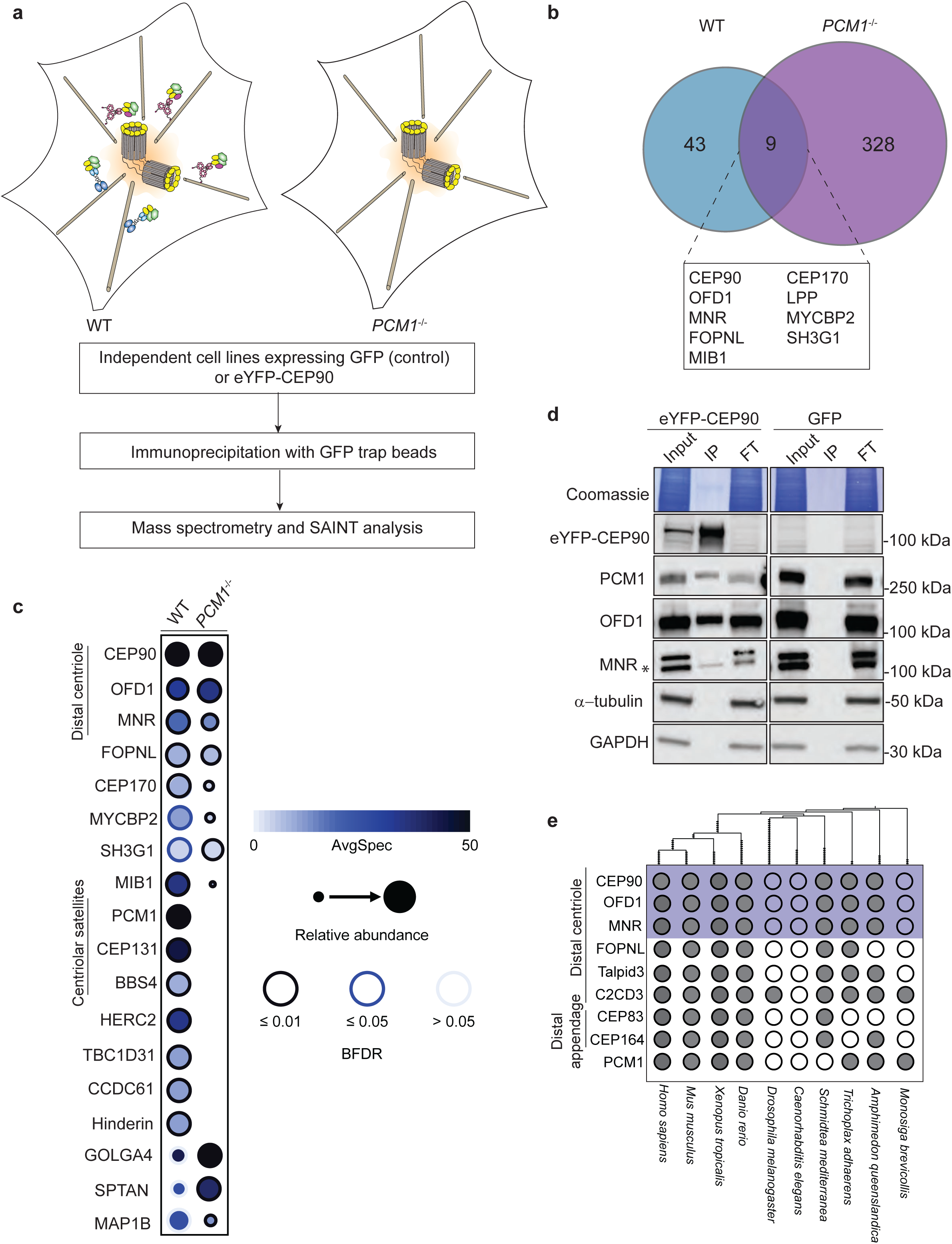
CEP90 forms a complex with OFD1 and MNR. **a.** Schematic depicting workflow used to identify CEP90 interactors at centriolar satellites and centrioles using WT and *PCM1*^-/-^ RPE1 cells. *PCM1*^-/-^ cells localize CEP90 to the distal centriole and WT cells localize CEP90 to the distal centriole and centriolar satellites. **b**. Venn diagram comparing high confidence interactors of CEP90 in WT and *PCM1*^-/-^ RPE1 cells. **c**. Interactome dot representation of selected CEP90 interactors identified in the proteomic screen. Hits were grouped based on their cellular localization. Average number of peptide spectra is represented by dot shade. Abundance of a peptide spectrum produced in relation to the most abundant spectrum is depicted by dot size. Bayesian false discovery rate (BFDR) is represented by rim color. **d**. Immunoblot of a subset of CEP90 interactions identified by proteomics were validated by co-immunoprecipitation (IP). CEP90 interacts with PCM1, OFD1 and MNR, but not α-tubulin or GAPDH. FT, flow through. Specific MNR band is indicated with an asterisk. The top band is nonspecific as it is undiminished in the MNR knockout cell lysates. **e.** Coulson plot showing the phylogenetic distribution of a subset of centriolar proteins in select ciliated metazoan species. Orthologs identified with high confidence are indicated with a filled circle, and a subset of CEP90 interactors further explored in this study are highlighted in blue. The dendrogram on top (made using iTOL (Ciccarelli et al., 2006)) shows the evolutionary relationship between species.

To identify centriolar interactors of CEP90, we assessed which interactors were detected in both WT and *PCM1*^-/-^ RPE1 cells, as these interactions are predicted to be independent of centriolar satellites. Notably, OFD1 and MNR, previously described components of centriolar satellites and centrosomes that interact with each other (Chevrier et al., 2016), were detected as CEP90 interactors (Fig. 2b and c). Using co-immunoprecipitation, we confirmed that CEP90 interacts with PCM1, MNR and OFD1 (Fig. 2d). Phylogenetic analysis revealed that CEP90, MNR and OFD1 are largely co-conserved in metazoans, but absent in ecdysozoa such as *Drosophila melanogaster* and *Caenorhabditis elegans* (Fig. 2e).

To assess whether MNR and OFD1 co-localize with CEP90, we immunostained RPE1 cells. CEP90 co-localized with MNR at centriolar satellites (Fig. 3a), and in cells treated with nocodazole to disperse centriolar satellites, at centrioles (Fig. 3b). Similarly, MNR and OFD1 co-localized at centriolar satellites (Fig. 3c), and centrioles in nocodazole treated cells (Fig. 3d). Thus, like CEP90 and OFD1, MNR is a component of both centriolar satellites and centrioles.

**Figure 3.**
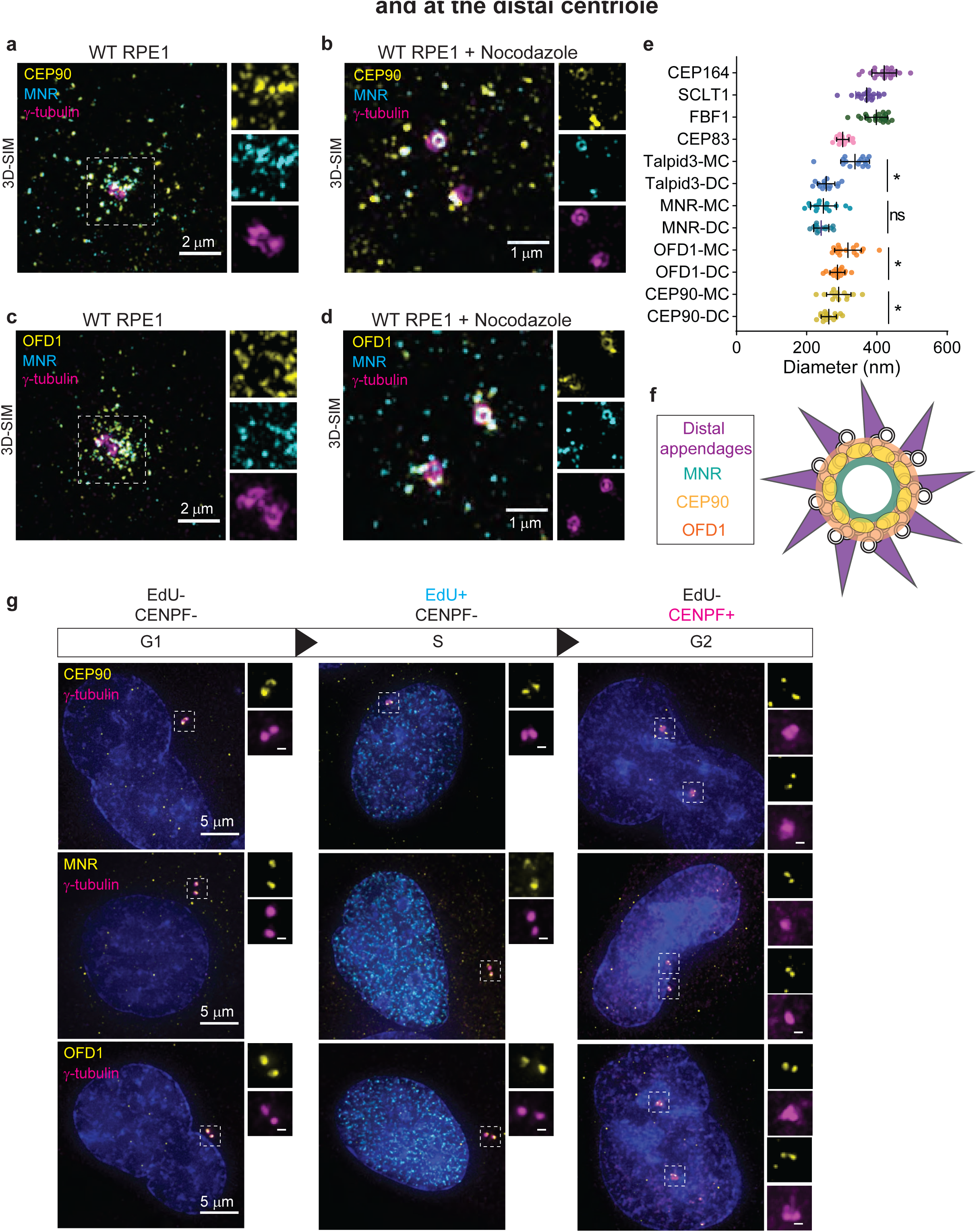
CEP90 co-localizes with OFD1 and MNR. **a.** Immunostaining of RPE1 cells for CEP90 (yellow), MNR (cyan) and γ-tubulin (magenta) demonstrating that CEP90 co-localizes with MNR at centriolar satellites. Scale bar = 2 μm. **b.** 3D-SIM of RPE1 cells treated with nocodazole to disperse centriolar satellites highlights co-localization of CEP90 (yellow) and MNR (cyan) at centrioles (γ-tubulin, magenta). Scale bar = 1 μm. **c**. Immunostaining of OFD1 (yellow), MNR (cyan) and γ-tubulin (magenta) reveals CEP90 and MNR co-localization at centriolar satellites. Scale bar = 2 μm. **d**. 3D-SIM of immunostained cells treated with nocodazole reveals a ring of OFD1 (yellow) co-localizing with MNR (cyan) at centrioles (γ-tubulin, magenta). Scale bar =1 μm. **e**. Measurements of ring diameters measured in 3D-SIM images. For distal centriole proteins, ring diameters were measured at the mother (MC) and daughter centriole (DC). Scatter dot plot shows mean ± SD. Asterisks indicate p <0.05, and ns indicate not significant determined using unpaired t test. n = 13-22 centrioles. **f**. Schematic representation of a radial view of a mother centriole with distal appendages (purple), OFD1 (orange), CEP90 (yellow) and MNR (green). **g**. Localization of distal centriole proteins (yellow) to centrioles (γ-tubulin, magenta) is cell cycle dependent. Cells in G1 were identified as lacking both EdU and CENPF staining, cells in S as being positive for EdU but lacking CENPF, and cells in G2 as being positive for CENPF but lacking EdU staining. Two puncta of CEP90, OFD1 and MNR localized were observed at one centrosome during G1 and S, and four puncta at two centrosomes after centrosome duplication during G2. Scale bar of main panel and insets = 5 μm and 0.5 μm respectively.

To gain insight into the spatial organization of distal centriole proteins with respect to distal appendages, we analyzed radially oriented centrioles with 3D-SIM (Fig. 3e). Consistent with previous observations, distal appendage components CEP164 and CEP83 were organized into respectively larger and smaller rings at the mother centriole (Yang et al., 2018; Bowler et al., 2019). Of the distal centriole proteins, MNR was organized into the smallest ring (244 ± 29 nm) (Fig. 3e and 3f). Talpid3, OFD1 and CEP90 also were organized into rings that, intriguingly, were of different diameters at the mother and daughter centrioles. At the mother centriole, rings, from largest to smallest, were comprised of Talpid3 (345 ± 27 nm), OFD1 (317 ± 37 nm) and CEP90 (291 ± 35 nm). At the daughter centriole, these rings were smaller and ordered differently from largest to smallest as OFD1 (288 ± 21 nm), CEP90 (263 ± 21 nm) and Talpid3 (255 ± 24 nm) (Fig. 3e and 3f), indicating that the distal centriole is reorganized when the daughter centriole matures into a mother.

Centriole biogenesis is coordinated with the cell cycle. Procentrioles form during S phase and elongate in G2. To ascertain when distal centriole proteins are recruited to centrioles, we used EdU incorporation and CENPF staining to distinguish between cells in different stages of the cell cycle (Viol et al., 2020). CEP90, OFD1 and MNR localized to mother and daughter centrioles in cells in G1 (EdU negative, CENPF negative) and in S phase (EdU positive, CENPF negative) (Fig. 3g). In G2 (EdU negative, CENPF positive), both centrosomes (marked by γ-tubulin) each possessed two puncta of CEP90, OFD1 and MNR (Fig. 3g). Therefore, CEP90, OFD1 and MNR are recruited to the distal end of elongating procentrioles in G2.

### CEP90 and MNR are critical for vertebrate development, ciliogenesis and Hedgehog signaling

To assess the function of CEP90 and MNR, we generated RPE1 cell lines lacking CEP90 and MNR using CRISPR/Cas9. Immunoblot analyses confirmed loss of protein in the mutant cell lines (Supplementary Fig 1 a-d). Notably, *CEP90*^-/-^ and *MNR*^-/-^ RPE1 cells possess centrioles but both lacked cilia (Fig. 4a and 4b). As we previously identified a role for OFD1 in cilium assembly (Singla et al., 2010; Hunkapiller et al., 2011), we conclude that the CEP90, MNR, OFD1 distal centriole complex is essential for ciliogenesis.

**Figure 4.**
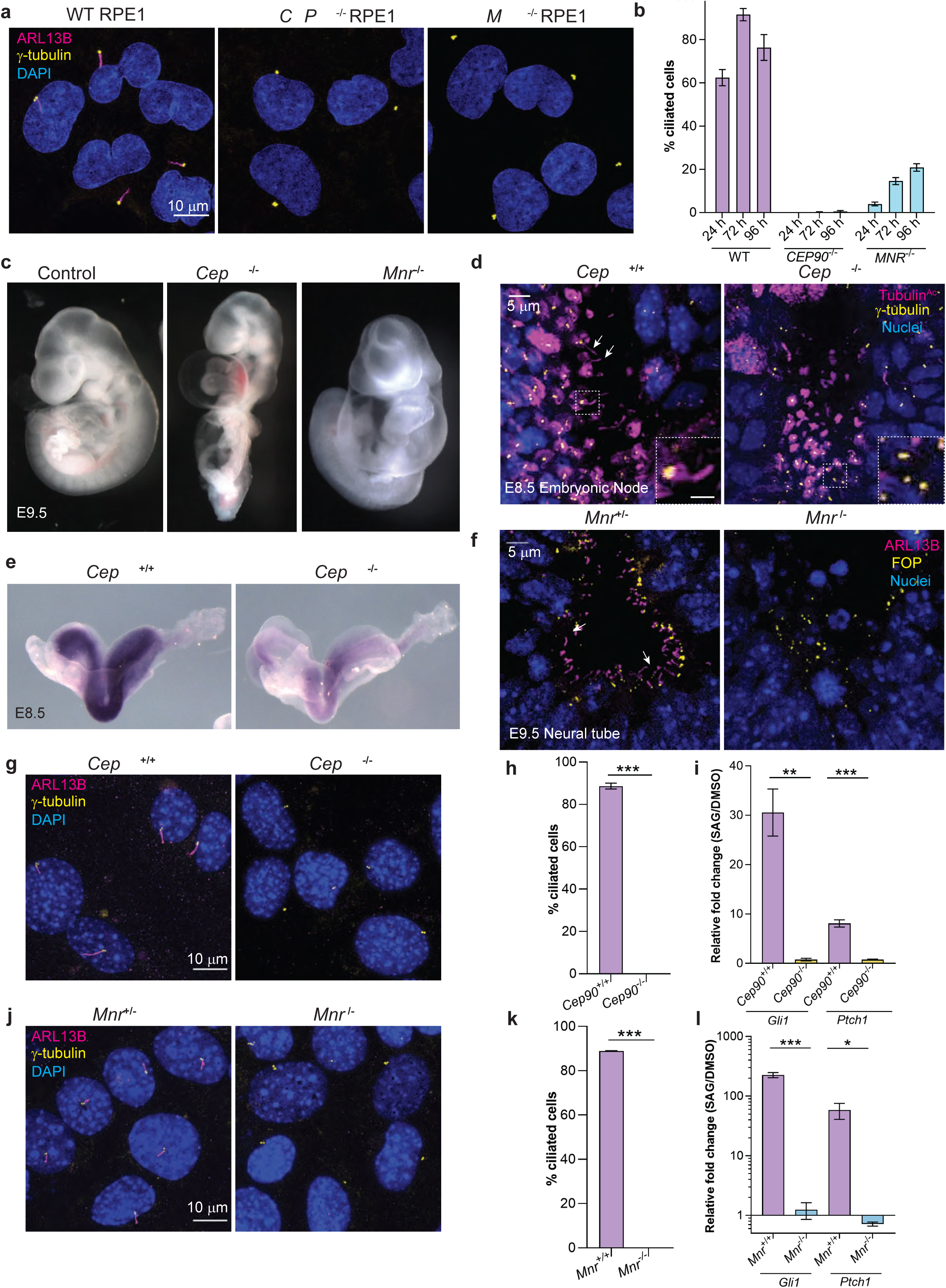
CEP90 and MNR are essential for ciliogenesis. **a**. WT, *CEP90*^-/-^ and *MNR*^-/-^ serum starved RPE1 cells were immunostained for cilia (ARL13B, magenta), centrosomes (γ-tubulin, yellow) and nuclei (Hoechst, blue). Scale bar = 10 μm. **b**. Quantification of ciliation frequency of WT, *CEP90*^-/-^ and *MNR*^-/-^ RPE1 cells serum starved for times indicated. n >100 cells from two biological replicates. **c.** Images of control, *Cep90*^-/-^ and *Mnr*^-/-^ embryos at embryonic day (E) 9.5. Arrows point to pericardial edema observed in mutant embryos. **d**. Whole-mount immunostaining of nodes of littermate control and *Cep90*^-/-^ embryos at E8.5 for centrosomes (γ-tubulin, yellow), cilia (Tubulin^Ac^, magenta) and nuclei (Hoechst, blue). Arrows in control image point to cilia projecting into the node lumen, which are absent in *Cep90*^-/-^ embryos. Scale bar of main panel and insets = 5 μm and 2 μm respectively. **e***. In situ* hybridization for *Gli1* in E8.5 littermate control and *Cep90*^-/-^ embryos, revealing decreased expression in the absence of CEP90 indicative of disrupted Hedgehog signaling. Scale bar = 5 μm. **f**. Immunostaining of E9.5 neural tube sections of littermate control and *Mnr*^-/-^ embryos for centrosomes (FOP, yellow), cilia (ARL13B, magenta) and nuclei (Hoechst, blue), indicating that MNR is required for ciliogenesis in vivo. Scale bar = 5 μm. **g**. MEFs derived from *Cep90*^+/+^ and *Cep90*^-/-^ embryos were serum starved for 24 h and immunostained for cilia (ARL13B, magenta), centrosomes (γ-tubulin, yellow) and nuclei (Hoechst, blue). Scale bar = 10 μm. **h**. Quantification of ciliation frequency shows loss of cilia in *Cep90*^-/-^ MEFs. Bar graph shows mean ± SEM. Asterisks indicate p <0.05 determined using unpaired t test. n >100 cells from two biological replicates. **i**. qRT-PCR of HH target genes *Gli1* and *Ptch1* in serum-starved *Cep90*^+/+^ and *Cep90*^-/-^ MEFs stimulated with 200nM SAG for 24h relative to DMSO-treated controls. Bar graph shows mean ± SEM. Asterisks indicate p <0.05 determined using unpaired t test. n = 3 biological replicates. **j**. MEFs derived from *Mnr*^+/+^ and *Mnr*^-/-^ embryos were serum starved for 24 h and immunostained for cilia (ARL13B, magenta), centrosomes (γ-tubulin, yellow) and nuclei (Hoechst, blue). Scale bar = 10 μm. **k**. Quantification of ciliation frequency shows loss of cilia in *Mnr*^-/-^ MEFs. Bar graph shows mean ± SEM. Asterisks indicate p <0.05 determined using unpaired t test. n >100 cells from two biological replicates. **l**. qRT-PCR of HH target genes *Gli1* and *Ptch1* in serum-starved *Mnr*^+/+^ and *Mnr*^-/-^ MEFs stimulated with 200nM SAG for 24h relative to DMSO-treated controls. Bar graph shows mean ± SEM. Asterisks indicate p <0.05 determined using unpaired t test. n = 3 biological replicates.

To query the function of CEP90 and MNR in vertebrates, we obtained *Cep90* and *Mnr* mutant mice from IMPC. The *Cep90* mutant mice (*Pibf1*^tm1.1 (KOMP)Vlcg^) contain a deletion of 7 exons, and the *Mnr* mutant mice (*4933427D14Rik*^tm1 (KOMP)Vlcg^) contain a deletion of 10 exons.

Heterozygous *Cep90* and *Mnr* mice were viable and fertile, with no obvious phenotypes. Homozygous *Cep90* and *Mnr* mice did not survive beyond embryonic day (E) 9.5. At E9.5, *Cep90*^-/-^ and *Mnr*^-/-^ embryos displayed pericardial edema (Fig. 4c) and unlooped, midline hearts. One key role for primary cilia is in transducing extracellular signals including Hedgehog proteins, secreted morphogens required for vertebrate development. As lack of Smoothened, a central component of the Hedgehog signal transduction pathway, or lack of cilia also produces unlooped, midline hearts (Corbit et al., 2005; Huangfu et al., 2003), we examined the presence of cilia in *Cep90*^-/-^ and *Mnr*^-/-^ embryos. γ-tubulin-positive centrioles were detected in both wild type and *Cep90*^-/-^ embryonic nodes at E8.5, but cilia were absent from *Cep90*^-/-^ nodes (Fig. 4d). Consistent with a defect in ciliogenesis, *Cep90*^-/-^ embryos also displayed attenuated expression of the Hedgehog target gene, *Gli1*, indicating defects in Hedgehog signal transduction (Fig. 4e). Similarly, cilia detected by ARL13B staining were absent in neural tube sections from E9.5 *Mnr*^-/-^ embryos (Fig. 4f). Mouse embryonic fibroblasts (MEFs) derived from *Cep90*^-/-^ and *Mnr*^-/-^ embryos recapitulated the ciliogenesis phenotype observed in *CEP90*^-/-^ and *MNR*^-/-^ RPE1 cells (Fig. 4g, h, j, k). Furthermore, qRT-PCR analysis confirmed that induction of Hedgehog target genes, *Gli1* and *Ptch1,* in response to the Hedgehog pathway agonist SAG was abrogated in *Cep90*^-/-^ and *Mnr*^-/-^ MEFs (Fig. 4 i, l). Thus, CEP90 and MNR are both required for vertebrate development, ciliogenesis, and Hedgehog signaling.

### CEP90 ciliopathy mutations affect ciliogenesis and centriolar satellite morphology

To identify regions in CEP90 required for cilium assembly, we tested whether CEP90 lacking either the N-terminus, C-terminus or central domain (Kim et al., 2012) could support ciliogenesis in *CEP90*^-/-^ RPE1 cells (Fig. 5a). Unlike expression of the full length CEP90 (FL-CEP90), CEP90 lacking the N terminal (CEP90^363-757^) or the C-terminal (CEP90^1-363^) regions failed to rescue ciliogenesis defects in *CEP90*^-/-^ RPE1 cells (Fig. 5b,c). A construct of CEP90 lacking residues required for interaction with PCM1 (CEP90^Δ271-363^) localized to the centrosomal region, and partially rescued ciliogenesis defects in *CEP90*^-/-^ RPE1 cells (Fig. 5a-c). These data support a role for both N and C-terminal regions of CEP90 in ciliogenesis, further highlighting its function as a critical centriolar scaffold.

**Figure 5.**
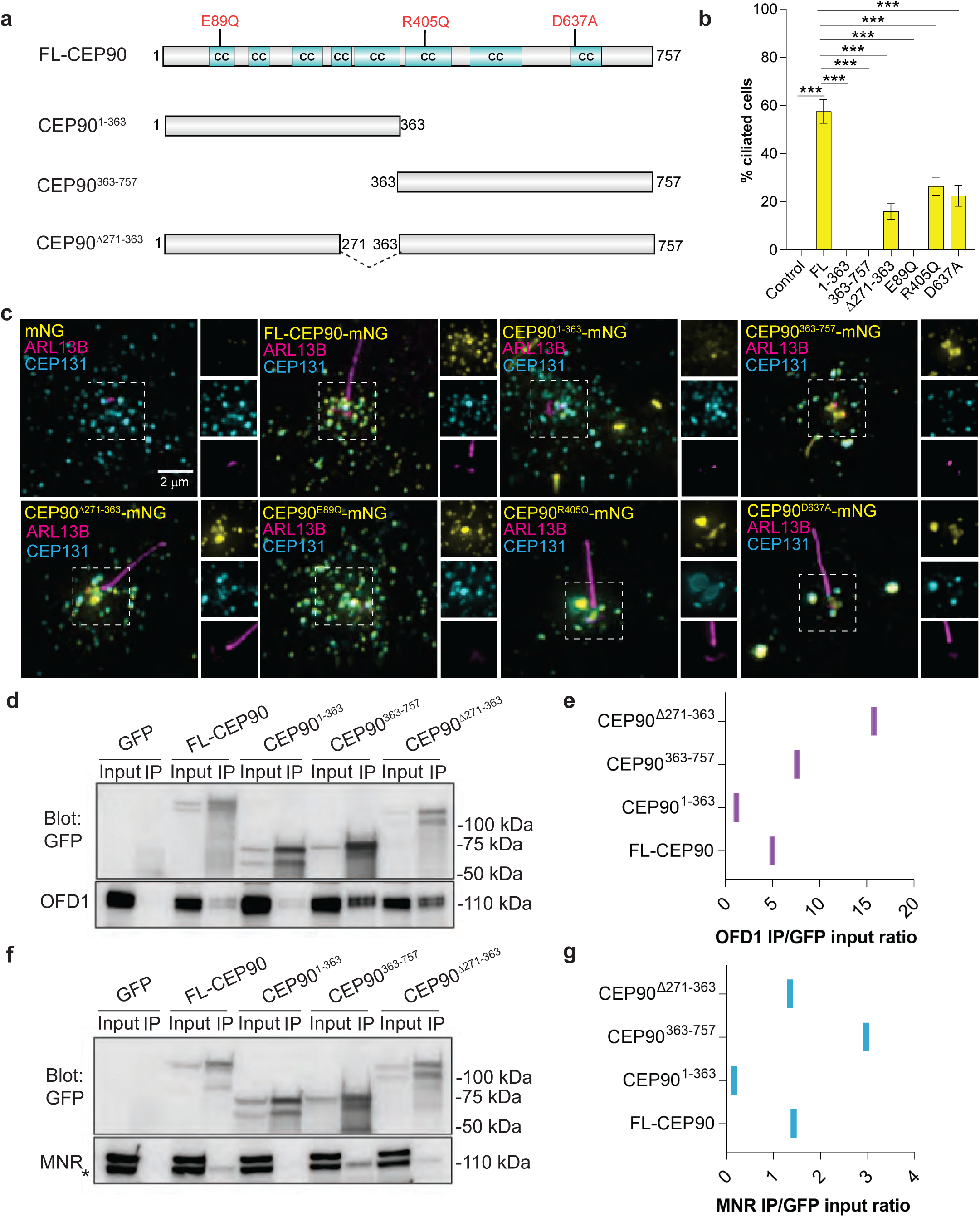
*CEP90* ciliopathy mutations affect ciliogenesis and centriolar satellite morphology. **a**. Schematic representation of full length (FL) human CEP90, truncation constructs and disease-associated mutations. Coiled-coil (CC) domains identified by MARCOIL (Zimmermann et al., 2018) at 90% threshold are indicated in cyan. **b**. Quantification of ciliation frequency of *CEP90*^-/-^ RPE1 (control), and *CEP90*^-/-^ cells expressing mNeonGreen (mNG) tagged disease-variants and truncations of CEP90. Graph shows mean ± SEM, and asterisks indicate p <0.05 determined by one-way ANOVA. n >100 cells from two biological replicates. **c**. *CEP90*^-/-^ RPE1 cells, and *CEP90*^-/-^ cells expressing the denoted mNG-tagged versions of CEP90 were serum starved for 24 h and immunostained for cilia (ARL13B), centriolar satellites (CEP131) and mNeonGreen. Scale bar = 2 μm. **d**. Immunoprecipitation of GFP-tagged FL and truncation constructs of CEP90 blotted for GFP and OFD1. **e**. Quantification of OFD1 band intensities relative to corresponding GFP input band intensities. **f**. Immunoprecipitation of GFP-tagged FL and truncation constructs of CEP90 blotted for GFP and MNR. **g**. Quantification of MNR band intensities relative to corresponding GFP input band intensities. Asterisk represents MNR band, top band is non-specific.

We also examined whether disease-associated variants of CEP90 compromise its ability to support cilia assembly or centriolar satellite localization. Previously, we identified a rare variant in CEP90 (CEP90^E89Q^) associated with microcephaly (Kodani et al., 2015). Surprisingly, this variant of CEP90 displayed normal localization to centrioles and centriolar satellites (Fig. 5c), but failed to support ciliogenesis in *CEP90*^-/-^ RPE1 cells (Fig. 5b,c). We also assessed two mutant forms of CEP90 (CEP90^R405Q^ and CEP90^D637A^) implicated in Joubert syndrome (Wheway et al., 2015) (Fig. 5b,c). Interestingly, both Joubert associated variants of CEP90 perturbed centriolar satellite morphology (Fig. 5c), and showed reduced ability to support ciliogenesis in *CEP90*^-/-^ RPE1 (Fig. 5b). Thus, our data suggest that distinct human disease-associated mutations in *CEP90* differentially compromise centriolar satellite morphogenesis and ciliogenesis.

To identify regions of CEP90 required for interaction with DISCO components, OFD1 and MNR, we performed co-immunoprecipitation experiments with CEP90 truncation constructs. Both OFD1 and MNR preferentially interacted with the C-terminal region of CEP90 (CEP90^363-757^), CEP90 lacking the C-terminal (CEP90^1-363^) failed to immunoprecipitate OFD1 and MNR (Fig 5d-g). Therefore, the C-terminal region of CEP90 (in which mutations cause Joubert syndrome) is critical for interaction with DISCO subunits, OFD1 and MNR (Wheway et al., 2015).

### MNR, but not CEP90, restricts centriolar length

Since distal centriole proteins OFD1, Talpid3 and C2CD3 control centriole length, we analyzed *CEP90*^-/-^ and *MNR*^-/-^ centrioles labeled with Tubulin^Ac^ and distal centriole protein CEP162 by 3D-SIM. Compared to control centrioles, centriole length was not altered in *CEP90*^-/-^ cells. However, centriole length was highly variable in *MNR*^-/-^ cells, with ∼30% of *MNR*^-/-^ cells containing hyper-elongated centrioles (some of which were over a micron long, Fig 6. a-c). The hyper-elongated centrioles in *MNR*^-/-^ cells possessed the distal centriole component CEP162 at one end, suggesting that proximal-distal polarity is maintained despite the elongation (Fig. 6 a). Serial section transmission electron microscopy (TEM) analysis confirmed the presence of elongated centrioles in *MNR*^-/-^ RPE1 cells (Fig. 6 d-e). Both mother and daughter centrioles (distinguished by localization of Ninein to the mother centriole-specific sub-distal appendage) were hyper-elongated in *MNR*^-/-^ cells (Fig. 6f). Therefore, MNR restrains centriole lengthening of both mother and daughter centrioles (Fig. 6f-g).

**Figure 6.**
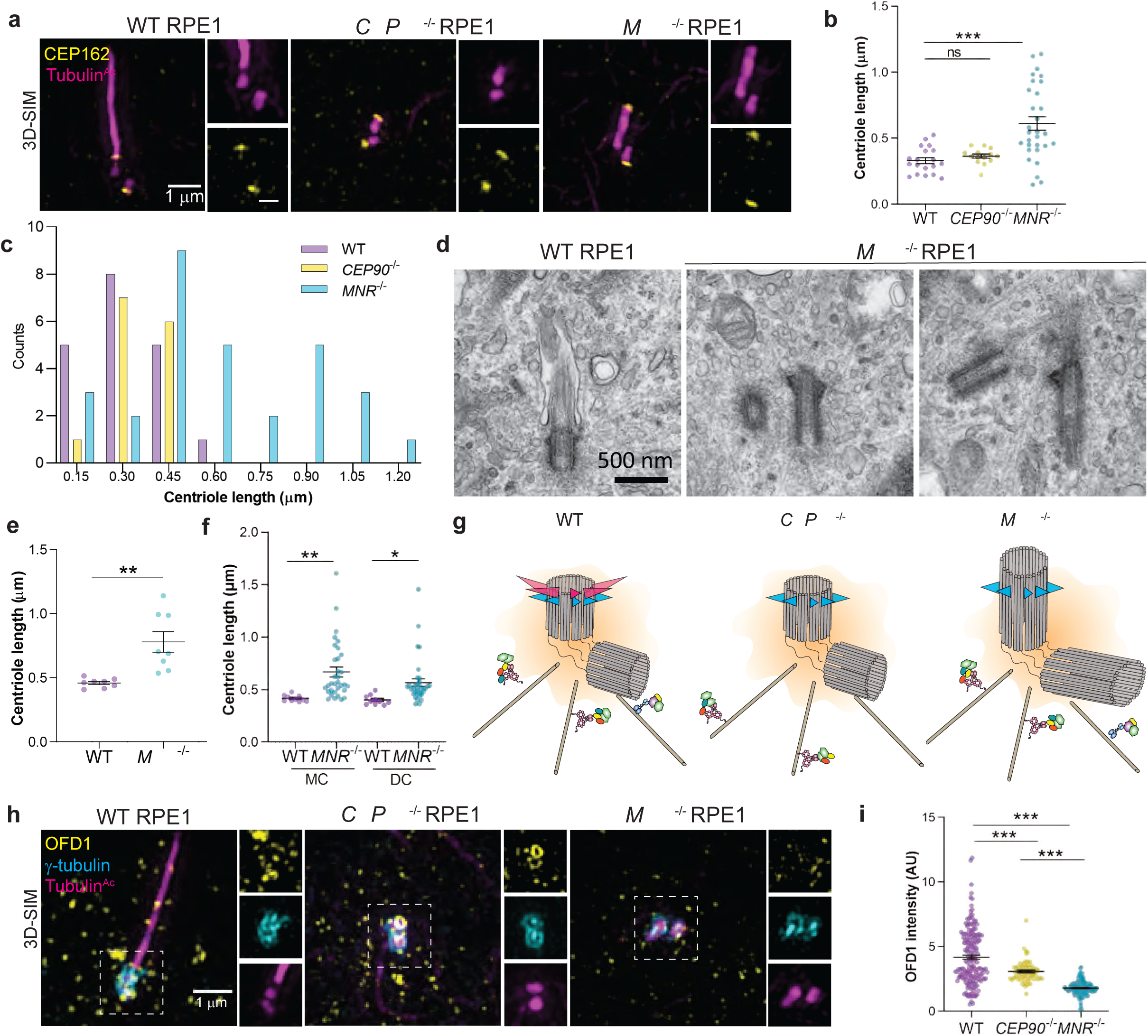
MNR, but not CEP90, restricts centriole length. **a.** 3D-SIM images of WT, *CEP90*^-/-^ and *MNR*^-/-^ RPE1 cells immunostained for cilia/centrioles (Tubulin^Ac^, magenta) and CEP162 (yellow), a distal centriolar protein. Scale bar of main panel and insets = 1 μm and 0.5 μm respectively.**b**. Graph of centriolar lengths measured using 3D-SIM images. Centriole lengths are longer and have a wider distribution in *MNR*^-/-^ cells than in WT or *CEP90*^-/-^ cells. Lines indicate mean ± SEM. Asterisks indicate p <0.05 determined using one-way ANOVA. n = 14-30 centrioles. **c.** Histogram of centriole lengths observed in WT, *MNR*^-/-^ and *CEP90*^-/-^ cells. n = 14-30 centrioles. **d.** Serial section TEM confirms the presence of elongated centrioles in *MNR*^-/-^ RPE1 cells. Scale bar = 500 nm. **e.** Centriole lengths of WT and *MNR*^-/-^ RPE1 cells measured using TEM images. Horizontal lines indicate means ± SEM, and asterisks indicate p<0.05 determined using unpaired t test. n = 8 centrioles per condition. **f**. WT and *MNR*^-/-^ RPE1 cells were serum-starved and immunostained with antibodies to Tubulin^Ac^ and sub-distal appendage component Ninein to distinguish mother (MC) and daughter centrioles (DC). Graph of centriolar lengths measured using 3D-SIM. MNR restrains centriole lengthening of both mother and daughter centrioles. Lines indicate mean ± SEM. Asterisks indicate p <0.05 determined using unpaired t-test. n = 10-35 measurements. **g**. Schematic depicting distinct roles of CEP90 and MNR in regulating centriole length. **h**. 3D-SIM imaging of serum-starved WT, *CEP90*^-/-^ and *MNR*^-/-^ RPE1 cells immunostained for OFD1 (yellow), centrioles (γ-tubulin, cyan) and cilia (Tubulin^Ac^, magenta). Boxed regions are depicted in insets throughout. OFD1 localizes to centrioles in WT and *CEP90*^-/-^, but not *MNR*^-/-^ cells. Scale bar = 1 μm. **i**. Quantification of OFD1 fluorescence intensity at centrioles in WT, *CEP90*^-/-^ and *MNR*^-/-^ cells. Horizontal lines indicate means ± SEM. Asterisks indicate p<0.05 determined using one-way ANOVA. n = 64-187 cells.

The abnormal centriole morphology in *MNR*^-/-^ cells is reminiscent of the phenotype we previously observed in *OFD1* mutant cells (Singla et al., 2010). Therefore, we examined the localization of OFD1 in *MNR*^-/-^ RPE1 cells. OFD1 failed to localize to the centrioles in the absence of MNR, but not CEP90 (Fig. 6 h-i). Overexpressed MNR localizes to microtubules (Chevrier et al., 2016) and microtubule-associated MNR can sequester endogenous OFD1 (Supplementary figure 2). Therefore, MNR is necessary and sufficient to recruit OFD1 (Fig. 6h-i), and our data support a model in which MNR recruits OFD1 to restrict centriole length(Feng et al., 2017; Srivastava and Panda, 2017), with CEP90 being dispensable for centriole length control and dedicated to ciliogenesis.

### CEP90 and MNR are required for the removal of CP110 and CEP97 from the mother centriole at the initiation of ciliogenesis

Ciliogenesis depends on a specialized protein transport machinery called intraflagellar transport (IFT). We examined whether IFT88 localization to basal bodies depends on CEP90 or MNR, and found that IFT88 was decreased at the mother centriole of both *CEP90*^-/-^ and *MNR*^-/-^ RPE1 cells in both serum-starved (Fig. 7 a,b) and cycling cells (Supplementary Fig. 3a-b).

**Figure 7.**
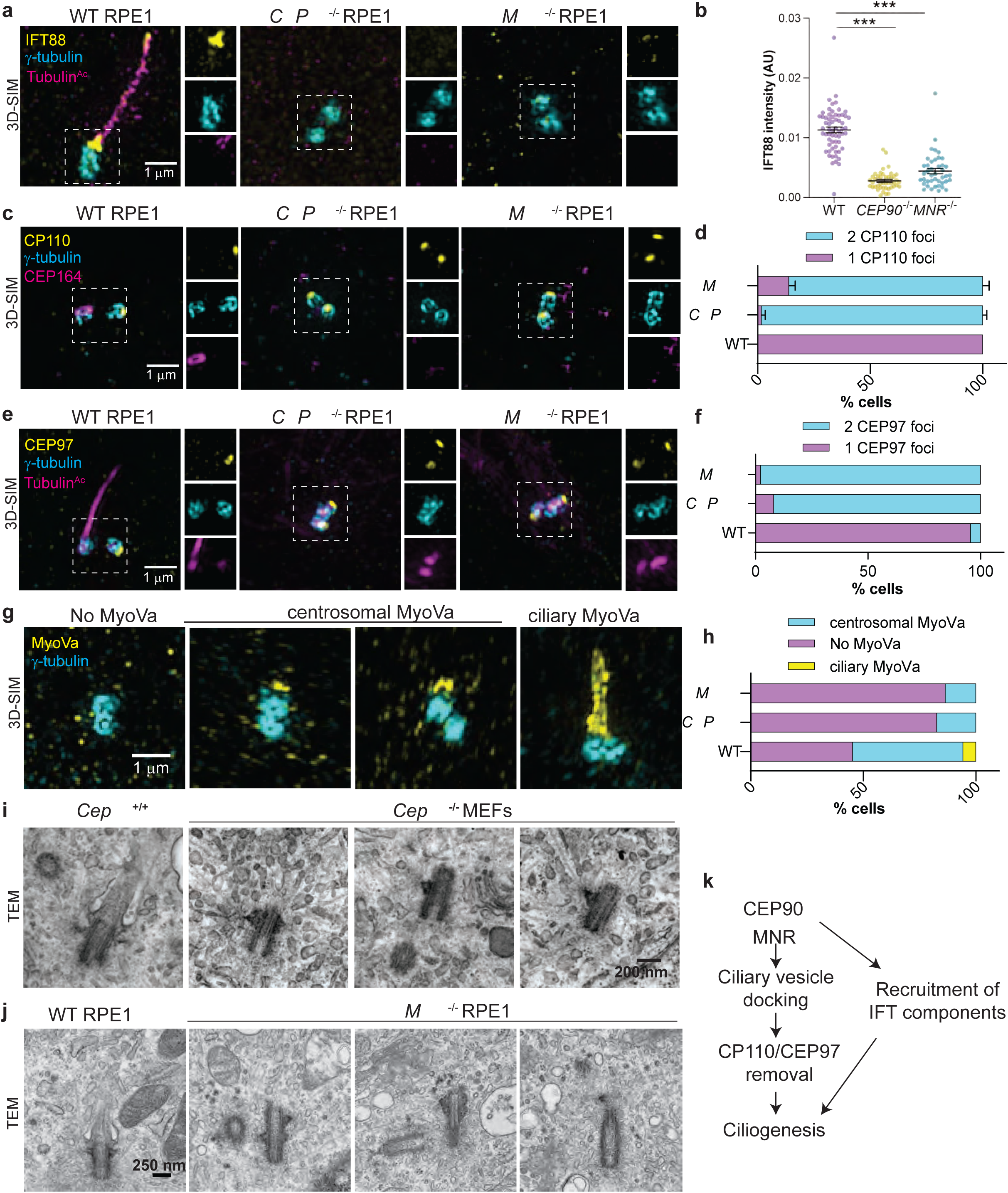
CEP90 and MNR are required for early ciliogenesis. **a** WT, *CEP90*^-/-^ and *MNR*^-/-^ RPE1 cells immunostained for IFT88 (yellow), centrioles (γ-tubulin, cyan) and cilia (Tubulin^Ac^, magenta. IFT88 is not recruited to the centrosome of *CEP90*^-/-^ or *MNR*^-/-^ cells. **b.** Quantification of IFT88 fluorescence intensity at WT, *CEP90*^-/-^ and *MNR*^-/-^ centrosomes. Lines indicate mean ± SEM. Asterisks indicate p <0.05 determined using unpaired t test. n = 38-63 measurements. **c**. WT, *CEP90*^-/-^ and *MNR*^-/-^ serum-starved RPE1 cells immunostained for CP110 (yellow), centrioles (γ-tubulin, cyan) and cilia (Tubulin^Ac^, magenta). Scale bar = 1 μm. **d**. Quantification of whether CP110 localizes to one or two centrioles. In the absence of CEP90 or MNR, CP110 continues to localize to the distal mother centriole. n>50 cells from 2 independent experiments. Scale bar = 1 μm. **e**. WT, *CEP90*^-/-^ and *MNR*^-/-^ serum-starved RPE1 cells immunostained for CEP97 (yellow), γ-tubulin (cyan) and Tubulin^Ac^ (magenta). Scale bar = 1 μm. **f**. Quantification of whether CEP97 localizes to one or two centrioles. As with CP110, CEP90 and MNR are required to remove CEP97 from the distal mother centriole. n>50 cells from 2 independent experiments. Scale bar = 1 μm. **g**. 3D-SIM images of RPE1 cells immunostained for Myo-Va (yellow) and centrioles (γ-tubulin, magenta). Myo-Va can not localize near centrosomes (left), can localize to preciliary vesicles (denoted centrosomal Myo-Va, middle), or can localize to the ciliary pocket (denoted ciliary Myo-Va, right). Scale bar = 1 μm. **h**. Quantification of three distinct Myo-Va staining patterns in WT, *CEP90*^-/-^ and *MNR*^-/-^ cells. n>50 cells from 2 independent experiments. **i.** Serial section TEM images of serum-starved *Cep90*^+/+^ and *Cep90*^-/-^ MEFs confirms the absence of preciliary vesicle docking at the *Cep90*^-/-^ mother centriole. Scale bar = 200 nm. n= 9 cells for both genotypes. **j.** Serial-section TEM images of WT and *MNR*^-/-^ RPE1 cells confirms the absence of preciliary vesicle vesicle docking at the *MNR*^-/-^ mother centriole. Scale bar = 250 nm. n= 10 cells for WT and n= 6 for *MNR*^-/-^ cells. **k**. Model based on our data highlighting the role of CEP90 and MNR in maturation of the mother centriole.

A critical early step in ciliogenesis is the removal from the distal mother centriole of two proteins that can inhibit ciliogenesis, CP110 and CEP97. We tested whether CEP90 or MNR function removal of CP110 and CEP97 by examining *CEP90*^-/-^ and *MNR*^-/-^ RPE1 cells. We found that cells lacking either CEP90 or MNR fail to remove CP110 and CEP97 (Fig. 7c-f). Taken together, these results reveal that CEP90 and MNR are required for early steps of ciliogenesis.

### CEP90 and MNR are required for ciliary vesicle docking and distal appendage assembly

As ciliary vesicle formation contributes to removing CP110 and CEP97 from the mother centriole (Westlake et al., 2011; Lu et al., 2015), we investigated whether persistence of CP110 and CEP97 in *CEP90*^-/-^ and *MNR*^-/-^ cells was due to defective recruitment of preciliary vesicles.

To assess whether CEP90 and MNR affect preciliary vesicle recruitment, we examined the localization of Myosin-Va-positive preciliary vesicles (Wu et al., 2018) to centrioles in wild type, *CEP90*^-/-^ and *MNR*^-/-^ RPE1 cells. Consistent with the defect in distal appendage formation, cells lacking either CEP90 or MNR showed reduced Myosin-Va at the mother centriole (Fig. 7g-h). To confirm the requirement for both CEP90 and MNR in preciliary vesicle recruitment, we examined wild-type and knockout cells by serial section TEM, which confirmed that centrioles in cells lacking CEP90 or MNR fail to dock to preciliary vesicles (Fig. 7i-j and Supplementary Figures 4). Thus, CEP90 and MNR are essential for ciliary vesicle recruitment to the mother centriole, a key early step of ciliogenesis (Fig. 7k).

An early step of ciliogenesis is the acquisition of appendages by the mother centriole, defining its maturation into a basal body. Sub-distal appendages mediate the anchoring of the basal body to microtubules and regulate the spatial positioning of the cilium in the cell (Mazo et al., 2016). In *CEP90*^-/-^ and *MNR*^-/-^ RPE1 cells, the localization of components of sub-distal appendages, such as Ninein and CEP170, were unaffected (Supplementary Fig 5a-d), indicating that CEP90 and MNR are dispensable for subdistal appendage formation.

A function of distal appendages is the recruitment of small preciliary vesicles to the mother centriole which fuse and give rise to the ciliary membrane (Tanos et al., 2013; Schmidt et al., 2012a). *CEP90*^-/-^ and *MNR*^-/-^ centrioles fail to recruit distal appendage proteins CEP83, FBF1, SCLT1, ANKRD26 and CEP164, indicating that they are essential for distal appendage formation (Fig. 8c-g). FBF1 and CEP164 failed to localize to the mother centriole in cycling *CEP90*^-/-^ and *MNR*^-/-^ cells, indicating that CEP90 and MNR organize the distal appendage irrespective of whether serum starvation initiates ciliogenesis (Supplementary Fig. 3e-h). CEP90 and MNR are not critical for distal appendage protein abundance (Supplementary Fig 6e), suggesting that they are not required for distal appendage protein stability but rather are essential for distal appendage assembly at the mother centriole.

**Figure 8.**
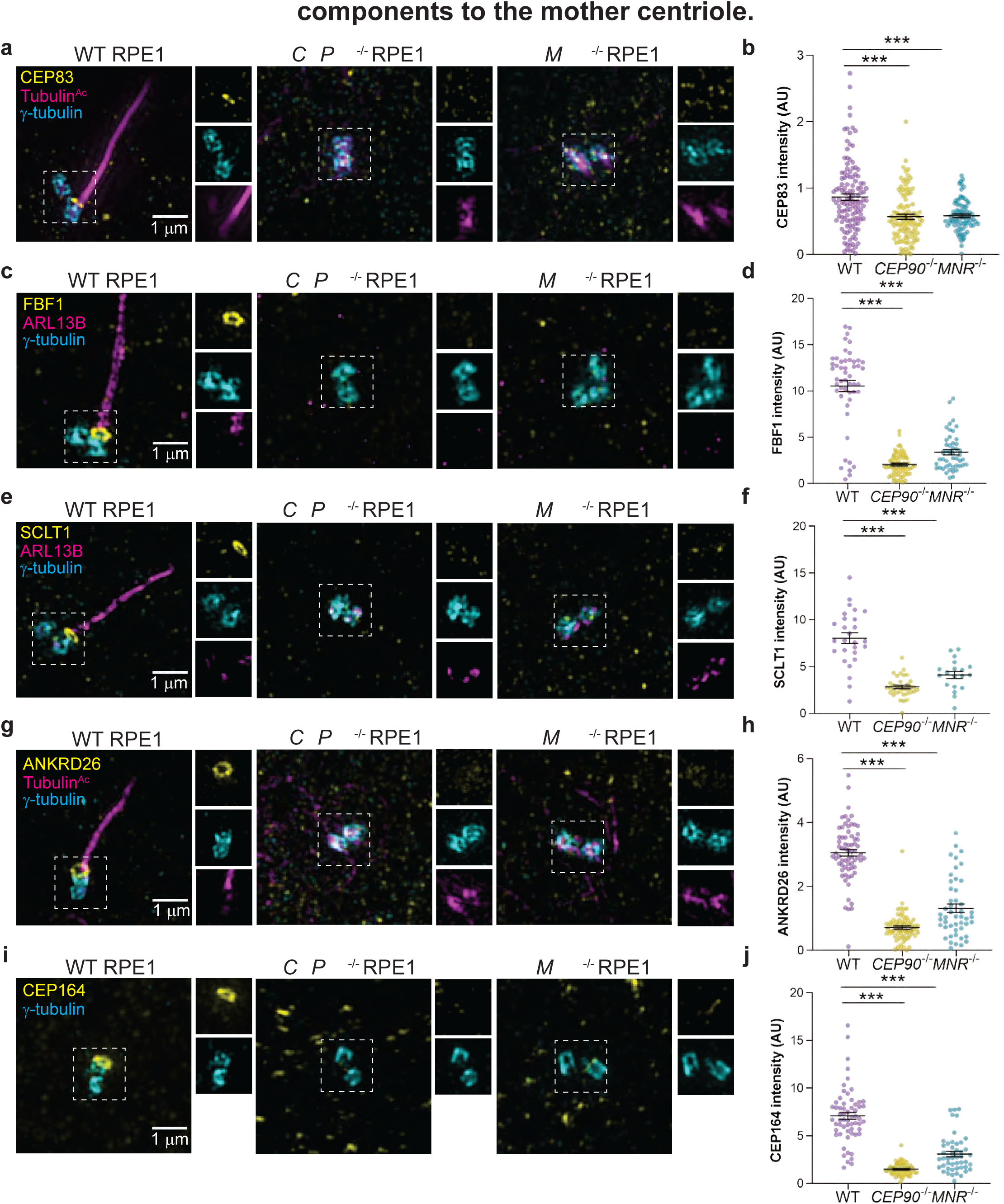
CEP90 and MNR recruit distal appendage components to the mother centriole. 3D-SIM images and quantification of centrosomal intensity of WT, *CEP90*^-/-^ and *MNR*^-/-^ RPE1 cells immunostained for γ-tubulin (cyan), Tubulin^Ac^ (magenta) and distal centriole components (yellow) CEP83 (**a,b**), FBF1 (**c,d**), SCLT1 (**e,f**), ANKRD26 (**g,h**) and CEP164 (**i,j**). Scale bar = 1 μm. Horizontal lines in scatter dot plots indicate means ± SEM. Asterisks indicate p<0.001 determined by one-way ANOVA. n ≥ 40 measurements per condition.

Recent studies show that daughter centriole proteins are removed from the mother centriole to enable distal appendage assembly (Wang et al., 2018; Mahjoub et al., 2010). Since CEP90 localizes to the distal daughter centriole as well as the distal mother centriole, we hypothesized that CEP90 may interact with distal daughter centriole proteins. Co-immunoprecipitation revealed that CEP90 did interact with daughter centriole components Centrobin and CEP120 (Supplementary Fig. 5f). As removing daughter centriole proteins requires distal centriole proteins, such as Talpid3, we examined the localization of Talpid3 and found that Talpid3 is recruited to the distal centriole in both wild-type and *CEP90*^-/-^ and *MNR*^-/-^ RPE1 cells (Supplementary Fig. 5g and h). Similarly, Centrobin and CEP120 localized to only one centriole in both wild-type and *CEP90*^-/-^ and *MNR*^-/-^ RPE1 cells (Supplementary Fig. 5 i-l). Therefore, CEP90 and MNR regulate distal appendage formation through a mechanism independent of Talpid3 recruitment or daughter centriole protein removal.

### CEP90 functions at the mother centriole to support distal appendage formation

Using siRNA-mediated gene knockdown, a previous study showed that CEP90 promotes centriolar satellite accumulation in the vicinity of centrosomes. We confirmed that, in *CEP90*^-/-^ RPE1 cells, PCM1-positive centriolar satellites failed to accumulate around centrosomes (Fig 9a and 9b). Interestingly, loss of MNR did not affect targeting of PCM1-positive centriolar satellites to the centrosome (Fig 9a and 9b), indicating that CEP90 and MNR have some distinct functions in centriolar satellite distribution. These observations raised the interesting possibility that CEP90-dependent targeting of centriolar satellites to the centrosomal area promotes distal appendage formation and ciliogenesis.

**Figure 9.**
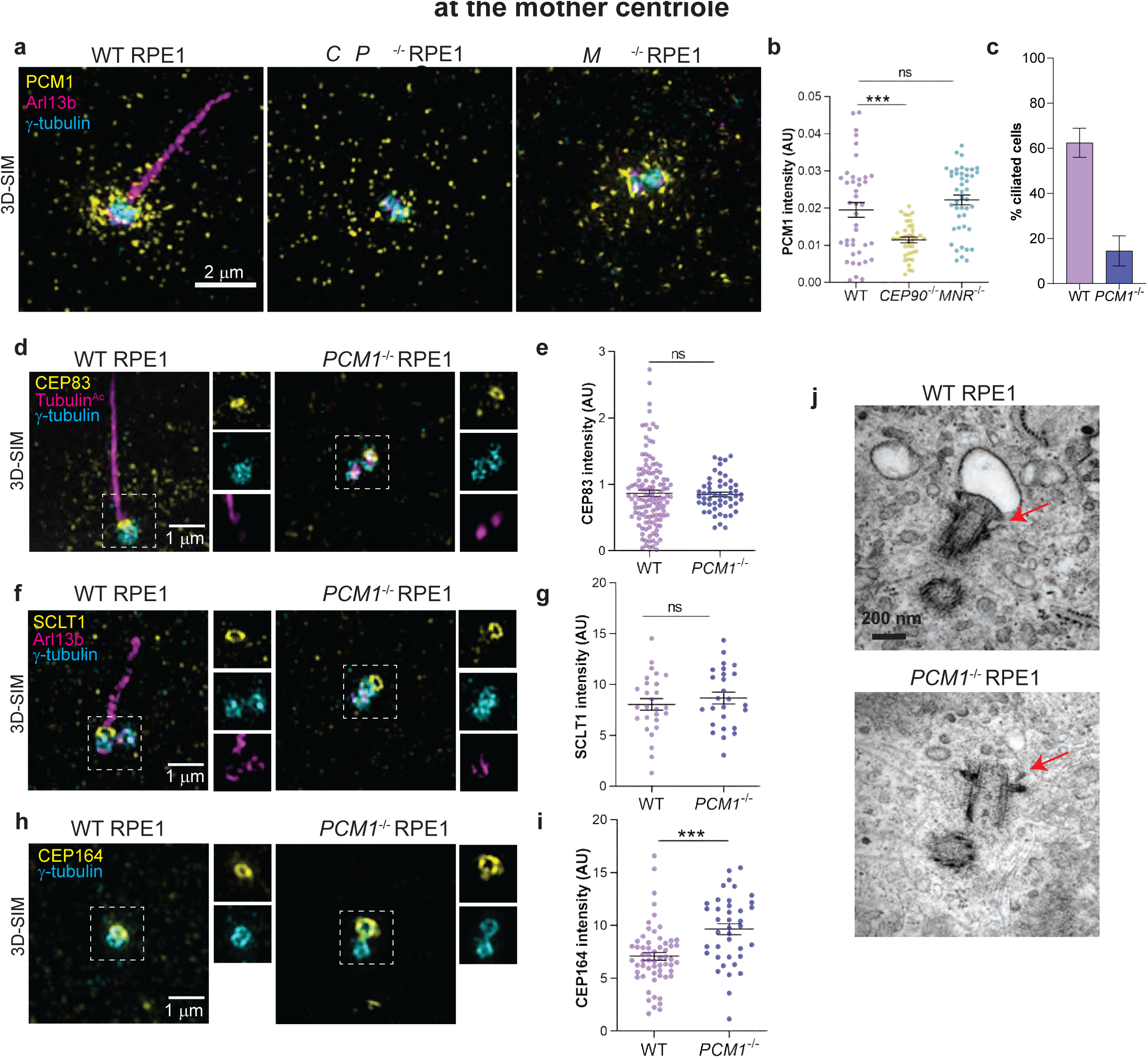
Centriolar satellites are dispensable for distal appendage assembly at the mother centriole. **a.** 3D-SIM images of WT, *CEP90*^-/-^ and *MNR*^-/-^ RPE1 cells immunostained with antibodies to PCM1, a centriolar satellite marker, ARL13B and γ-tubulin. Scale bar = 2 μm. **b**. Quantification of PCM1 intensity in the pericentrosomal area. PCM1-positive centriolar satellites fail to accumulate around centrosomes in *CEP90*^-/-^, but not in *MNR*^-/-^ cells. Lines indicate mean ± SEM. Asterisks indicate p <0.05 determined using ordinary one-way ANOVA. n = 37-44 measurements. **c**. Ciliogenesis is disrupted in *PCM1*^-/-^ RPE1 cells at 24 h post serum starvation. n >100 cells from two biological replicates. **d**. WT and *PCM1*^-/-^ RPE1 cells were serum starved for 24 h and stained with antibodies to CEP83, γ-tubulin (centrosome marker) and Tubulin^Ac^ (cilia marker). 3D-SIM imaging reveals ring of CEP83 at the mother centrioles in *PCM1*^-/-^ RPE1 cells. Scale bar = 1 μm. **e**. Quantification of CEP83 fluorescence intensity at centrioles. Scatter dot plots show mean ± SEM. NS = not significant, determined using unpaired t test. n = 53-124 measurements. **f**. WT and *PCM1*^-/-^ RPE1 cells were serum starved for 24 h and stained with antibodies to SCLT1, γ-tubulin (centrosome marker) and ARL13B (cilia marker). 3D-SIM imaging reveals ring of SCLT1 at the mother centrioles in WT and *PCM1*^-/-^ RPE1 cells. Scale bar = 1 μm. **g**. Quantification of SCLT1 fluorescence intensity at centrioles. Scatter dot plots show mean ± SEM. NS = not significant, determined using unpaired t test. n = 26-28 measurements. **h**. WT and *PCM1*^-/-^ RPE1 cells were serum starved for 24 h and stained with antibodies to CEP164 and γ-tubulin (centrosome marker). 3D-SIM imaging reveals ring of CEP164 at the mother centrioles in *PCM1*^-/-^ RPE1 cells. Scale bar = 1 μm. **i**. Quantification of CEP164 fluorescence intensity at centrioles. Scatter dot plots show mean ± SEM. Asterisks indicate p< 0.001, determined using unpaired t test. n = 40-60 measurements. **j**. Representative TEM images of WT and *PCM1*^-/-^ RPE1 cells serum starved for 1 hour. Distal appendages are marked with red arrows.

As *PCM1*^-/-^ cells lack centriolar satellites but retain CEP90 at centrioles (Fig. 1c), these cells can help disentangle the functions of CEP90 and MNR at centriolar satellites and centrioles. In accordance with previous observations (Wang et al., 2016; Odabasi et al., 2019), RPE1 cells lacking PCM1 displayed compromised ciliogenesis (Fig 9c). In stark contrast to *CEP90*^-/-^ cells, distal appendage components CEP83, SCLT1and CEP164 localized equivalently to the mother centrioles of *PCM1*^-/-^ and wild type RPE1 cells (Fig 9d-i), indicating that distal appendage assembly is independent of PCM1. Furthermore, serial section transmission electron microscopy confirmed the presence of distal appendages in *PCM1*^-/-^ RPE1 cells (Fig 9j). As the critical centriolar satellite scaffold PCM1 is not required for either CEP90 localization to centrioles or distal appendage formation, we conclude that the centriolar satellite population of CEP90 is dispensable for distal appendage assembly.

If the centriolar satellite population is dispensable, we predicted that MNR and OFD1 recruitment of CEP90 to the distal centriole would be critical for distal appendage assembly and ciliogenesis. Indeed, we found that although CEP90 protein levels were unchanged in *MNR*^-/-^ RPE1 cells (Supplementary Fig. 1d), CEP90 failed to localize to *MNR*^-/-^ centrioles, while MNR localized to the centrioles in the absence of CEP90 (Fig. 10 a-d).

**Figure 10.**
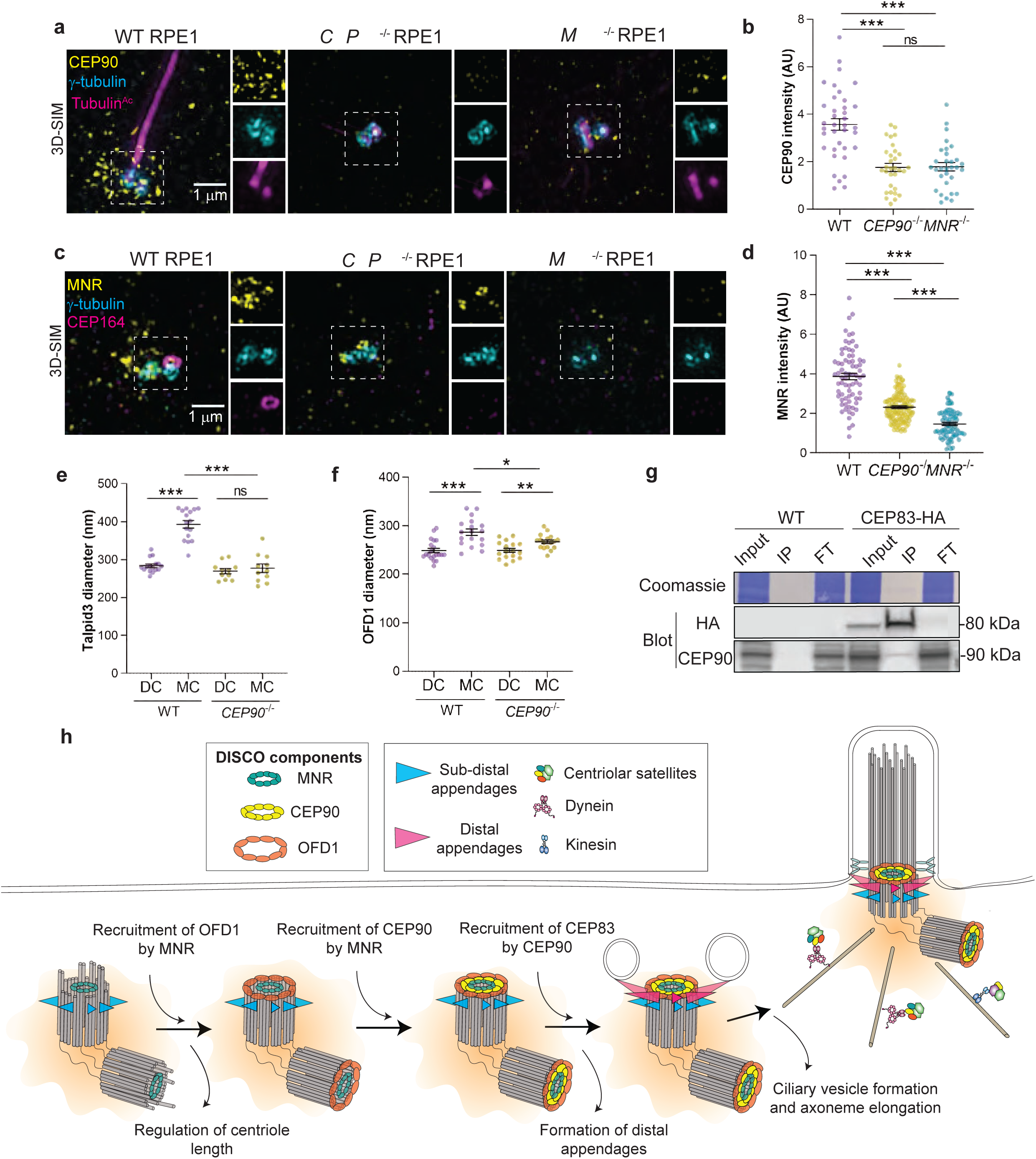
MNR recruits CEP90 which recruits CEP83 to build distal appendages. **a.** 3D-SIM imaging of WT, *CEP90*^-/-^ and *MNR*^-/-^ serum-starved RPE1 cells immunostained for CEP90 (yellow), γ-tubulin (cyan) and Tubulin^Ac^ (magenta). Scale bar = 1 μm. **b**. Quantification of CEP90 fluorescence intensity at centrioles. Horizontal lines in scatter dot plots indicate means ± SEM. Asterisks indicate p < 0.0001, ns = not significant, determined using one-way ANOVA. n = 31-37 measurements. CEP90 fails to localize to *MNR*^-/-^ and *CEP90*^-/-^ centrioles, although protein levels of CEP90 remained unchanged in *MNR*^-/-^ RPE1 cells (Supplementary Fig. 1d). **c**. 3D-SIM imaging of WT, *CEP90*^-/-^ and *MNR*^-/-^ serum-starved RPE1 cells immunostained for MNR (yellow), γ-tubulin (cyan) and CEP164 (magenta). Scale bar = 1 μm. **d.** Quantification of MNR fluorescence intensity at centrioles. Horizontal lines in scatter dot plots indicate means ± SEM. Asterisks indicate p < 0.0001, determined using one-way ANOVA. n = 73-122 measurements. MNR localization is reduced but present at *CEP90*^-/-^ centrioles. **e**. Quantification of Talpid3 diameter at mother (MC) and daughter centrioles (DC) in serum-starved WT and *CEP90*^-/-^ cells. Horizontal lines in scatter dot plots indicate means ± SEM. Asterisks indicate p < 0.05 determined using one-way ANOVA, ns = not significant. n = 11-16 measurements. The Talpid3 ring diameter is increased at WT mother centrioles, but not at *CEP90*^-/-^ mother centrioles. **f**. Quantification of OFD1 diameter at mother (MC) and daughter centrioles (DC) in serum-starved WT and *CEP90*^-/-^ cells. Horizontal lines in scatter dot plots indicate means ± SEM. Asterisks indicate p < 0.05, determined using one-way ANOVA. n = 17-20 measurements. In the absence of CEP90, the mother centriole ring of OFD1 does not expand as in wild-type cells. **g**. WT or RPE1 cells expressing CEP83-HA were lysed, immunoprecipitated with anti-HA antibody bound beads and immunoblotted with antibodies to HA and CEP90. **h**. Model of the hierarchical recruitment of the DISCO complex. MNR recruitment of OFD1 restrains centriole elongation and MNR recruits CEP90 which, in turn, recruits CEP83, the base of the distal appendage.

To investigate how CEP90 can be specifically required for the formation distal appendages, we investigated whether CEP90 reorganizes distal centriole proteins at the mother centriole to promote distal appendage assembly. 3D-SIM analysis revealed that Talpid3 and OFD1 diameters were smaller in mother centrioles of *CEP90*^-/-^ cells compared to WT (Fig 10e and 10f). These results support a model where CEP90 functions in the transformation of a daughter to a mother centriole by reorganizing the distal centriole.

Next, we examined whether CEP90 interacts with the distal centriole base. Co-immunoprecipitation analysis revealed an interaction between CEP90 and the proximal most-component of the distal centriole, CEP83 (Fig. 10g). We propose that MNR recruits CEP90 to the distal centriole, and CEP90, in turn, recruits CEP83 to initiate distal appendage assembly (Fig. 10h). Therefore, distal centriolar proteins are recruited in a hierarchical manner, with MNR recruiting OFD1 to restrict centriole length, and CEP90 dedicated to distal appendage assembly.

## DISCUSSION

### CEP90 and MNR are critical for distal appendage assembly and ciliogenesis

Through the process of mitosis, vertebrate daughter cells each inherit two centrioles. Yet, only the older, mother centriole templates a cilium. In no small part, this unique function of the mother centriole depends on its distal appendages, acquired during the previous G2/M phase. In this study, we identify a multi-protein complex which we name DISCO, comprised of CEP90, OFD1 and MNR and critical for distal appendage formation and ciliogenesis.

Consequent to their role in distal appendage formation, CEP90 and MNR are also required for subsequent events in ciliogenesis, including recruitment of the preciliary vesicles that give rise to the ciliary membrane, and loading of IFT88 (Sillibourne et al., 2013; Tanos et al., 2013). Unlike some other centriolar proteins, such as CEP120 (Tsai et al., 2019), CEP90 and MNR are specifically required for distal appendage formation, but dispensable for subdistal appendage formation, revealing that discrete mother centriolar complexes support distal and subdistal appendage assembly.

In mice, we found that CEP90 and MNR are essential for embryonic development, and particularly for Hedgehog signaling. Primary cilia are required for Hedgehog signal transduction in vertebrates. A central component of the Hedgehog pathway is the seven-pass transmembrane protein, Smoothened (Smo). Interestingly, *Cep90, Mnr* and *Smo* mutants arrested at similar points in embryonic development and displayed similar phenotypes, emphasizing the importance of CEP90 and MNR in ciliary Hedgehog signaling.

### MNR recruits OFD1 to regulate centriole length

CEP90 and MNR colocalize with their interactor OFD1 at the distal ends of centrioles. We found that distal centriole proteins are recruited to the centriole in a hierarchical manner. MNR forms the innermost ring at the distal centriole and is essential for the recruitment of OFD1 and CEP90, whereas CEP90 is dispensable for the recruitment of the other complex members. Overexpressed MNR localizes to microtubules (Chevrier et al., 2016) bringing along OFD1. Therefore, we propose that MNR binds to the distal centriolar microtubules to recruit OFD1, culminating in CEP90 recruitment (Fig. 10).

We had previously demonstrated that OFD1 was essential to restrain centriolar length (Singla et al., 2010). We have found that one of its partners, MNR, is also critical to restrict centriolar elongation. However, its other partner, CEP90, does not control centriolar length. Thus, within the complex comprised of MNR, OFD1 and CEP90, the subunits have distinct functions, with CEP90 dedicated to founding distal appendages. Consequently, the requirement for MNR and OFD1 in distal appendage assembly may be secondary to their roles in recruiting CEP90 to the distal centriole. The most parsimonious model is that MNR (and perhaps OFD1) at the distal centriole recruits CEP90 to build distal appendages, and that MNR recruits OFD1 to restrict centriole elongation.

### CEP90 recruits CEP83 to initiate distal appendage assembly at the mother centriole

The functions of centriolar satellites, although intimately associated with centrosomes, remain unclear (Odabasi et al., 2019; Prosser and Pelletier, 2020). CEP90 is a component of centriolar satellites, where it interacts with PCM1, the major scaffolding protein of centriolar satellites (Kim et al., 2012). Like CEP90, its interactors MNR and OFD1 are centriolar satellite proteins. As loss of CEP90 disrupts the peri-centrosomal localization of centriolar satellites, we initially hypothesized that centriolar satellites transport CEP90 to centrioles, or that CEP90 acts at centriolar satellites to transport distal appendage components to the mother centriole. In *PCM1*^-/-^ RPE1 cells, centriolar satellites are disrupted but CEP90 remains at the distal centriole, indicating that satellites are dispensable for CEP90 localization at centrioles. Moreover, to our surprise, we found that *PCM1*^-/-^ RPE1 cells assemble distal appendages. Thus, centriolar satellites (and CEP90 at centriolar satellites) are dispensable for distal appendage assembly, indicating that CEP90 functions at the distal centriole to build distal appendages.

A combination of expansion and structured illumination super-resolution microscopy revealed that CEP90 decorates the distal centriole in a discontinuous ring-like pattern with nine-fold symmetry; this ring of CEP90 is proximal to and smaller than the ring of CEP164. While antibodies to endogenous CEP90 label the daughter centriole more heavily than the mother centriole, epitope-tagged CEP90 localizes equivalently to both the mother and daughter centrioles. We speculate that reduced accessibility of the CEP90 antibody once distal appendages are assembled at the mother centriole may limit endogenous CEP90 immunofluoresence.

At the centriole, CEP90 interacts with and is critical for the recruitment of the distal appendage component CEP83. As CEP83 is the root of the distal appendage (Tanos et al., 2013), CEP90 is required to initiate distal appendage formation. In support of this conclusion, no distal appendage-like structures were observable in serial section transmission electron micrographs of cells lacking CEP90. These data raise the tantalizing possibility that the nine-fold ring of CEP90 at the distal centriole templates the assembly of the distal appendages.

Super-resolved microscopy also revealed that OFD1 and Talpid3 are structurally different in the mother and daughter centrioles, with the diameter of both rings increasing upon transition from daughter to mother. CEP90 is essential for OFD1 and Talpid3 rings to dilate to the mother centriole-specific diameter. We propose that this CEP90-dependent reorganization of distal centriole directs adoption of mother centriole-specific properties, including distal appendage assembly and ciliogenesis. Future work will focus on establishing how CEP90 reorganizes OFD1 and Talpid3 at the distal mother centriole to initiate distal appendage assembly.

In addition to MNR, OFD1 and CEP90, Talpid3 and C2CD3 localize to the distal centriole and are required for distal appendage formation (Singla et al., 2010; Thauvin-Robinet et al., 2014; Ye et al., 2014; Wang et al., 2018). We did not identify C2CD3 or Talpid3 in our proteomic analysis, suggesting that they might be part of a distinct, distal centriole complex (Tsai et al., 2019). Although CEP90 interacts with daughter centriole proteins CEP120 and Centrobin, Talpid3 recruitment, and the subsequent removal Centrobin and CEP120 from the distal mother centriole occurs normally in *CEP90*^-/-^ RPE1 cells. Therefore, CEP90 acts independently or downstream of Talpid3 recruitment and daughter centriole protein removal to regulate distal appendage assembly.

Mutations in *CEP90* cause Joubert syndrome (Kodani et al., 2015; Wheway et al., 2015; Hebbar et al., 2018), while *OFD1* and *MNR* cause Orofaciodigital and Joubert syndrome (Singla et al., 2010; Stephen et al., 2017; Chevrier et al., 2016; Coene et al., 2009). Joubert and Orofaciodigital syndromes have partially overlapping clinical features. Ultrastructural studies have identified a cogwheel-like structure at the distal domain of human centrioles (Paintrand et al., 1992; Ibrahim et al., 2009), raising the possibility that DISCO may be components of this structure. We propose that DISCO both controls centriolar length and builds distal appendages, and that inherited defects in this cogwheel that compromise these critical functions attenuate ciliogenesis, Hedgehog signaling and embryonic patterning, resulting in Joubert and Orofaciodigital syndromes.

## MATERIAL AND METHODS

### Mouse lines

*Cep90* (*Pibf1*^tm1.1(KOMP)Vlcg^) and *Mnr* (*4933427D14Rik*^tm1.1(KOMP)Vlcg^) mice, generated in a C57BL/6NJ background were obtained from the International Mouse Phenotyping Consortium. Mice were housed in a barrier facility with veterinary supervision and given food and water *ad libitum*. All mouse protocols were approved by the Institutional Animal Care and Use Committee at the University of California, San Francisco and the University of Alabama, Birmingham.

### Cell lines and cell culture

Human retinal epithelial (RPE1-hTERT) cells were cultured in DMEM/F12 (Thermo Fisher; catalog 10565042) supplemented with 10 % FBS at 37 °C in 5% CO_2_. To induce ciliation, cells were serum starved in Opti-MEM reduced serum media for indicated times. 293T cells were cultured in DMEM supplemented with 10% FBS and 1X GlutaMAX at 37 °C in 5% CO_2_.

RPE1 cell lines stably expressing eYFP-CEP90 were generated using lentiviruses containing the human *CEP90* cDNA (gift from Kunsoo Rhee (Kim and Rhee, 2011; Kim et al., 2012)) in the pLVX-IRES-Puro (Clontech, Mountain View, CA) plasmid background. *CEP90*^-/-^ RPE1 cells expressing mNeonGreen-tagged full length, truncation and disease-associated mutation variants of CEP90 were generated using lentiviruses containing human *CEP90* cDNA in a pLVX-EF1α^Δ^-mNeonGreen plasmid background. Lentiviruses were generated using the Lenti-X Packaging Single Shot system (Takara Bio), according to manufacturer’s guidelines. Infected cells were plated in a glass-bottom 96 well plate (Cellvis; catalog P96-1.5H-N) using limiting dilution and monoclonal cell lines expressing eYFP-CEP90 were manually selected based on fluorescence.

RPE1 cells stably expressing HA-CEP83 under the control of tetracycline-inducible promoter were a gift from Barbara Tanos and Bryan Tsou (Tanos et al., 2013). CEP83 expression was induced with 1 μg/ml doxycycline (Fischer Chemical; Catalog BP26535) for 48 h.

*Cep90^-/-^* and *Mnr^-/-^* MEFs were derived from E8.5 embryos along with littermate WT control MEFs. Cells were cultured in DMEM supplemented with 10% FBS (Invitrogen) and Glutamax-I (Invitrogen), and subsequently immortalized by transduction with SV40 large T antigen.

All cell lines were routinely tested for mycoplasma contamination and found negative.

### Generation of *CEP90*^-/-^, *MNR*^-/-^ and *PCM1*^-/-^ RPE1 cells by CRISPR/Cas9 gene targeting

RNA guided targeting of *CEP90* and *MNR* was achieved by coexpression of Cas9 along with guide RNAs. The pSpCas9 (BB)-2A-GFP (PX458) was a gift from Feng Zhang (Addgene plasmid #48138; http://n2t.net/addgene:48138; RRID:Addgene_48138). The gRNA sequences used for CEP90 and MNR were 5′-GATGAGGAAATATCATCCGT-3′ and 5′-GTATAAAATACCCGACCACA-3′ respectively.

*PCM1*^-/-^ RPE1 cells were generated by electroporating recombinant Cas9 along with sgRNAs (CRISPRevolution sgRNA EZ kit, Synthego). The sgRNA targeting exon 4 of human *PCM1*: 5′-GAAAAGAAUAAGAAAAAGUU-3′ was used. 1.5 nmol sgRNA was resuspended in 15 μl nuclease-free TE for a final concentration of 100 μM (100 pmol/μl). The RNP mixture containing 1.8 μl of sgRNA with 3 μl (90 pmol) Truecut Cas9 v2 (Thermo Fisher; catalog A36498) in a total volume of 5 μl was incubated at room temperature for 15 minutes. The RNP mixture was electroporated into hTERT-RPE1 cells using the Neon transfection system (Thermo Fisher) according to manufacturer’s instruction using the following parameters: 1350 V pulse voltage, 20 ms pulse width and 2 pulses. All electroporated cells were serially diluted, and single colonies screened by western blotting, immunofluorescence and PCR analyses. For genotyping, the following PCR primers were used: 5′-TGGGATGCACTAAATTGCCTA-3′ and 5′-TTACCTGCCGTTTGAAGACA-3′ for *CEP90* alleles, 5′-TCCAGTGAACCAACTCACAGA-3′ and 5′-TAGGAGCGTGGCTGTGTCTAT-3′ for MNR alleles, and 5′-ACAGGCCATGTTAATTTTTGCT-3′ and 5′-CCATCCCCAGTGATTAAAATTC-3′ for *PCM1* alleles. PCR products were cloned and sequenced.

### Plasmids and transfections

Myc-DDK tagged *Mnr* (*4933427D14Rik*) cDNA in cloned in a pCMV6 plasmid was obtained from Origene (MR211309). hTERT-RPE1 cells were transfected using TransIT-LT1 transfection reagent (Mirus Bio) according to manufacturer guidelines.

For co-immunoprecipitation experiments in Fig. 5, C-terminal GFP-2x Strep-tagged full length and truncation versions of CEP90 were cloned into pLVX-EF1α-IRES-Puro backbone and transfected into 293T cells using TransIT-293 transfection reagent (Mirus Bio) according to manufacturer guidelines.

### Antibodies

The list of primary antibodies and the dilution they were used at are listed in Supplementary Table 1. Secondary antibodies conjugated to Alexa Fluor 488, 568 and 647 were purchased from Thermo Fisher Scientific, and used at a 1:500 dilution.

### Immunofluorescence

For immunostaining, cells were fixed with either 100% cold methanol for 3 minutes or 4% paraformaldehyde in dPBS for 15 minutes at room temperature, and then incubated in blocking buffer (2.5% BSA, 0.1% Triton-X 100 in PBS) for 1 hour at room temperature. Paraformaldehyde fixed cells were permeabilized with 0.1% Triton X-100 in PBS for 15 minutes at room temperature prior to addition of blocking buffer. Coverslips were incubated with primary antibodies in blocking buffer overnight at 4 °C, washed three times with PBS and incubated in secondary antibodies in blocking buffer for 1-2 h at room temperature. Nuclei were counterstained with Hoechst 33352 (Thermo Fisher Scientific; catalog H3570). Coverslips were washed three times with PBS and mounted with Prolong Diamond (Thermo Fisher Scientific; catalog P36961).

In experiments requiring disruption of the microtubule cytoskeleton, cells were treated with 20 μM Nocodazole (Sigma Aldrich; catalog SML1665) for 2 h at 37 °C prior to fixation.

Identification of cell cycle stages was performed as described previously (Viol et al., 2020), using the Click-iT EdU Alexa Fluor 555 imaging kit (Life Technologies). Briefly, *PCM1*^-/-^ RPE1 cells were treated with 10 μM EdU for 30 minutes and fixed in 100% cold methanol for 3 minutes. The Click-IT reaction was performed according to manufacturer guidelines, and samples subsequently processed for indirect immunofluorescence. Cells lacking CENPF and EdU in the nucleus were categorized as G1. EdU-positive, CENPF-negative cells were categorized as S, while cells with nuclear CENPF staining, without EdU staining were classified as being in G2.

### Super-resolution microscopy

3D-SIM and 2D-SIM was performed using the DeltaVision OMX-SR microscope (GE Healthcare) using the 60x/1.42 NA oil immersion PSF objective and three sCMOS cameras. Immersion oil with refractive index of 1.518 was used for most experiments. Z stacks of 5-6 μm were collected using a 0.125 μm step size. Raw images were reconstructed using SoftWorx 6.5.2 (GE Healthcare) using default parameters.

Label-retention expansion microscopy was performed as described previously (Shi et al., 2019). Briefly, RPE1 cells were cultured on 16-well chambered slides (Sigma; catalog GBL112358-8EA) coated with 0.1% gelatin (Sigma Aldrich; catalog G1393). Cells were serum starved in Opti-MEM to induce ciliation for 24 h prior to fixation. After incubation with primary antibodies, cells were incubated with appropriate secondary antibodies conjugated to NHS-MA-biotin or NHS-MA-DIG. After gel polymerization and proteinase K digestion, gels were stained with fluorescently labeled streptavidin or digitonin antibodies and images were acquired on the DeltaVision OMX-SR.

### Fluorescence intensity measurements and statistical analyses

For fluorescence intensity measurements, z stacks were acquired on the DeltaVision OMX-SR using widefield settings. Identical laser power and exposure settings were used for control and experimental samples. Average intensity projections were generated using Image J (Schneider et al., 2012), and images transferred to CellProfiler image analysis software (McQuin et al., 2018). Cilia number were quantified by using Hoechst stained nuclei to count the total number of cells, and a cilia marker (ARL13B/Tubulin^Ac^) to identify cilia using the object identification module in CellProfiler using difference in signal intensity and size to segment cilia. For quantification of centrosomal intensity, a mask around the centrosomal area was generated using a centrosomal marker (γ-tubulin/FOP) to identify the centrosome in CellProfiler. This centrosomal mask was used to determine fluorescence intensity (integrated intensity) and area (in pixels) for the channel of interest. Fluorescence intensity values in a pericentrosomal area were used to measure background. Cells were serum starved for 24 h prior to fixing, to synchronize cells in G0/G1 and eliminate cell cycle-dependent alterations in centrosomal proteins. Data were exported to Microsoft Excel, and graphs generated in GraphPad Prism 8.

Statistical analyses were performed using GraphPad Prism 8. Results represented are mean ± standard deviation. Statistical differences between data sets were analyzed using one-way Anova with Tukey’s multiple comparison tests or a two-tailed unpaired student’s t test. Data distribution was assumed to be normal but this was not formally tested. P value < 0.05 was considered significant and indicated with an asterisk. NS denotes not significant.

### Transmission electron microscopy

For electron microscopy, cells were plated on 8-well Permanox slides (Nunc; catalog177445), gently washed in 0.1 M phosphate buffer (PB) and fixed in 3.5% EM-grade glutaraldehyde (Electron Microscopy Sciences; catalog 16210) in 0.1M Phosphate buffer (PB) for 10 minutes at 37 °C. Fixative was removed, replaced with fresh fixative, and samples incubated at 4 °C for 1 h. Slides were washed three times with 0.1 M PB and processed for TEM as described previously (Singla et al., 2010).

### Co-immunoprecipitation and Western Blotting

Cells grown in 10 cm petri dishes were washed twice with ice-cold dPBS and lysed for 30 minutes on ice with frequent pipetting. Lysis buffer used was 10 mM Tris/Cl pH 7.5, 150 mM NaCl, 0.5 mM EDTA, 0.5 % Nonidet P40 substitute supplemented with 1x protease inhibitor (Roche cOmplete mini, EDTA free; catalog 4693159001) and 1x phosphatase inhibitor (Thermo Fisher PhosStop; catalog 4906845001) cocktails. Lysates were cleared by centrifugation at 17,000 g for 10 minutes at 4 °C. GFP-trap magnetic agarose (Chromotek; catalog gtma-10) was used for eYFP-CEP90 co-immunoprecipitation assays, and HA-magnetic beads (Pierce; catalog 88837) for CEP83-HA co-immunoprecipitation assays. Co-immunoprecipitation assays were performed according to manufacturer guidelines. Samples were separated on 4-15% Criterion TGX Precast gels (Bio-Rad), and transferred to Immobilon PVDF membranes (0.45 μm pore size) for chemiluminescence detection.

### Mass spectrometry and data analysis

Eluates obtained after co-immunoprecipitation were reduced by the addition of 1 mM DTT at 60°C for 15 minutes, cooled to room temperature, alkylated by the addition of 3 mM iodoacetamide for 45 minutes in the dark. Alkylation was quenched by the addition of 3 mM DTT and proteins were digested overnight at 37°C with 1 μg trypsin (0.5 μg/μl; Promega). Following digestion, peptides were acidified with trifluoroacetic acid (0.5% final, pH < 2), desalted using UltraMicroSpin Columns (PROTO 300 C18 300Å; The NEST Group) according to manufacturer’s specifications, and dried under vacuum centrifugation. Samples were resuspended in 4% formic acid, 4% acetonitrile solution, and separated by a reversed-phase gradient over a nanoflow column (360 μm O.D. x 75 μm I.D.) packed with 25 cm of 1.8 μm Reprosil C18 particles with (Dr. Maisch). The HPLC buffers were 0.1% formic acid and 100% acetonitrile on 0.1% formic acid for buffer A and B respectively. The gradient was operated at 400 nL/min from 0 to 28% buffer B over 40min, followed by a column wash at 95% B, with a total acquisition time of 50 min. Eluting peptides were analyzed in on a Bruke timsTOF Pro mass spectrometry system equipped with a Bruker nanoElute high-pressure liquid chromatography system interfaced via a captiveSpray source. A data-dependent PASEF acquisition (Meier et al., 2018) method was used for data acquisition using the following parameters: 100-1700 m/z range, 0.85-1.30 V*s/cm^2^ trapped ion mobility range, 1600V spray voltage, intensities of 200,000 were repeated for in PASEF 0 times, intensities of 100,000 to 200,000 were repeated for in PASEF 5 times, intensities of less than 100,000 were repeated for in PASEF 10 times, 4 PASEF MS/MS scans with a total cycle time of 0.53 s, and active exclusion for 0.4 min. Data was search against the human proteome database (canonical sequences downloaded from Uniprot 3/21/2018) using MaxQuant (Cox and Mann, 2008; Prianichnikov et al., 2020). Peptide and protein identifications were filtered to 1% false-discovery rate at the peptide and protein level, and protein-protein interaction analysis was performed using SAINTexpress (Teo et al., 2014). The mass spectrometry data files (raw and search results) have been deposited to the ProteomeXchange Consortium (Deutsch et al., 2017; Perez-Riverol et al., 2019) (http://proteomecentral.proteomexchange.org) via the PRIDE partner repository with dataset identifier PXD022372.

## Online supplemental material

**Fig. S1** depicts CRISPR/Cas9-mediated mutations of *CEP90*, *MNR* and *PCM1* and effect on corresponding proteins as assessed by immunoblotting. **Fig. S2** shows that overexpressed MNR localizes to microtubules and sequesters endogenous OFD1. **Fig. S3** shows that CEP90 and MNR regulate distal appendage assembly irrespective of whether the cell possesses a cilium. **Fig. S4** includes serial-section TEMs of WT, *Cep90*^-/-^ and *MNR*^-/-^ cells. **Fig. S5** shows that CEP90 and MNR regulate distal appendage assembly independent of Talpid3 recruitment and removal of daughter centriole proteins. **Table 1** lists all primary antibodies used in this study.

## ACKNOWLEDGEMENTS

We thank E Yu for help with animal husbandry and genotyping of mice used in this study. We thank members of the Reiter and Yoder labs for helpful discussions. This work was supported by NIH R01HD089918 to J.F.R and B.Y., NIH R01R01DE029454 and R01AR054396 to J.F.R., the Valencian Council for Innovation, Universities, Science and Digital Society (PROMETEO/2019/075) to J.M.G-V and the Spanish Ministry of Science, Innovation and Universities (PCI2018-093062) to V.H-P, NIH NIGMS R01GM124334 to B.H., by the NIH Pathway to Independence Award K99GM126136/R00GM126136 and the UCSF Mary Anne Koda-Kimble Seed Award for Innovation to X.S. D.K. is supported a Jane Coffin Childs Postdoctoral Fellowship, and the Program for Breakthrough Biomedical Research Award, which is partially funded by the Sandler Foundation. B. H. and J.F.R. are Chan Zuckerberg Biohub Investigators.

The authors declare no competing financial interests.

## Author contributions

Conceptualization: D. Kumar, B. Yoder and J.F Reiter; data acquisition: D. Kumar, A. Rains, V. Herranz-Perez, Q. Lu, X. Shi, D.L. Swaney, E. Stevenson; funding acquisition: D. Kumar, N. J. Krogan, C. Westlake, J. M. Garcia-Verdugo, B. Huang, B. Yoder and J.F. Reiter; writing - original draft: D. Kumar; writing - review and editing: D. Kumar, A. Rains, V. Herranz-Perez, B. Yoder, J.F. Reiter.

## SUPPLEMENTARY FIGURES

**SUPPLEMENTARY FIGURE 1.**
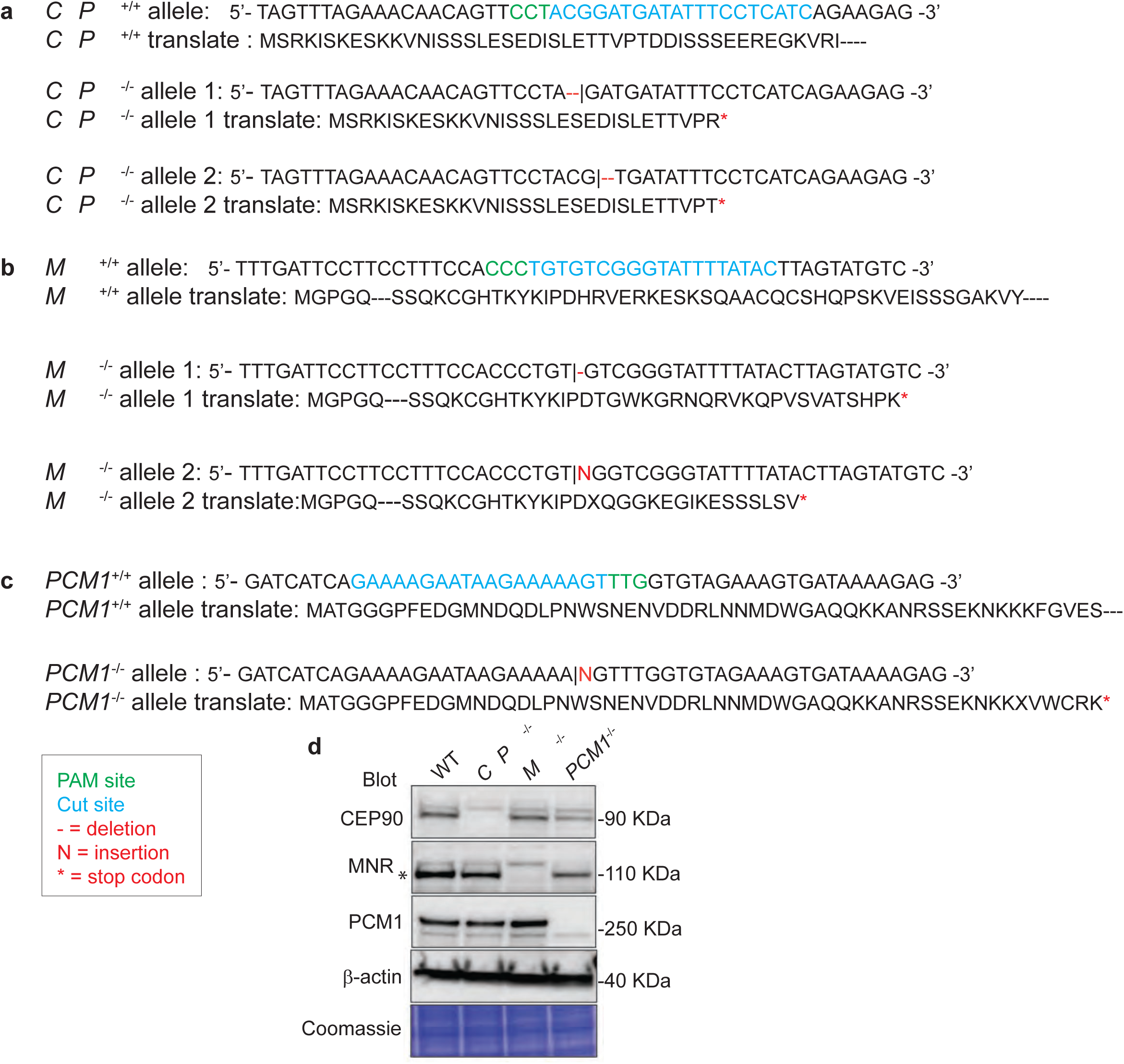
Generation of *CEP90*^-/-^, *MNR*^-/-^ and *PCM1*^-/-^ cell lines. Sequence analysis of genomic DNA isolated from control and *CEP90*^-/-^ (**a**), *MNR*^-/-^ (**b**) and *PCM1*^-/-^ (**c**) RPE1 cell lines generated using CRISPR-Cas9. Insertions and deletions, and translation products resulting from genome editing are indicated. **d**. Immunoblot analysis of whole cell lysates from control, *CEP90*^-/-^, *MNR*^-/-^ and *PCM1*^-/-^ RPE1 cell lines confirms loss of protein in mutant cell lines. Specific MNR band is indicated with an asterisk, top band is non-specific.

**SUPPLEMENTARY FIGURE 2.**
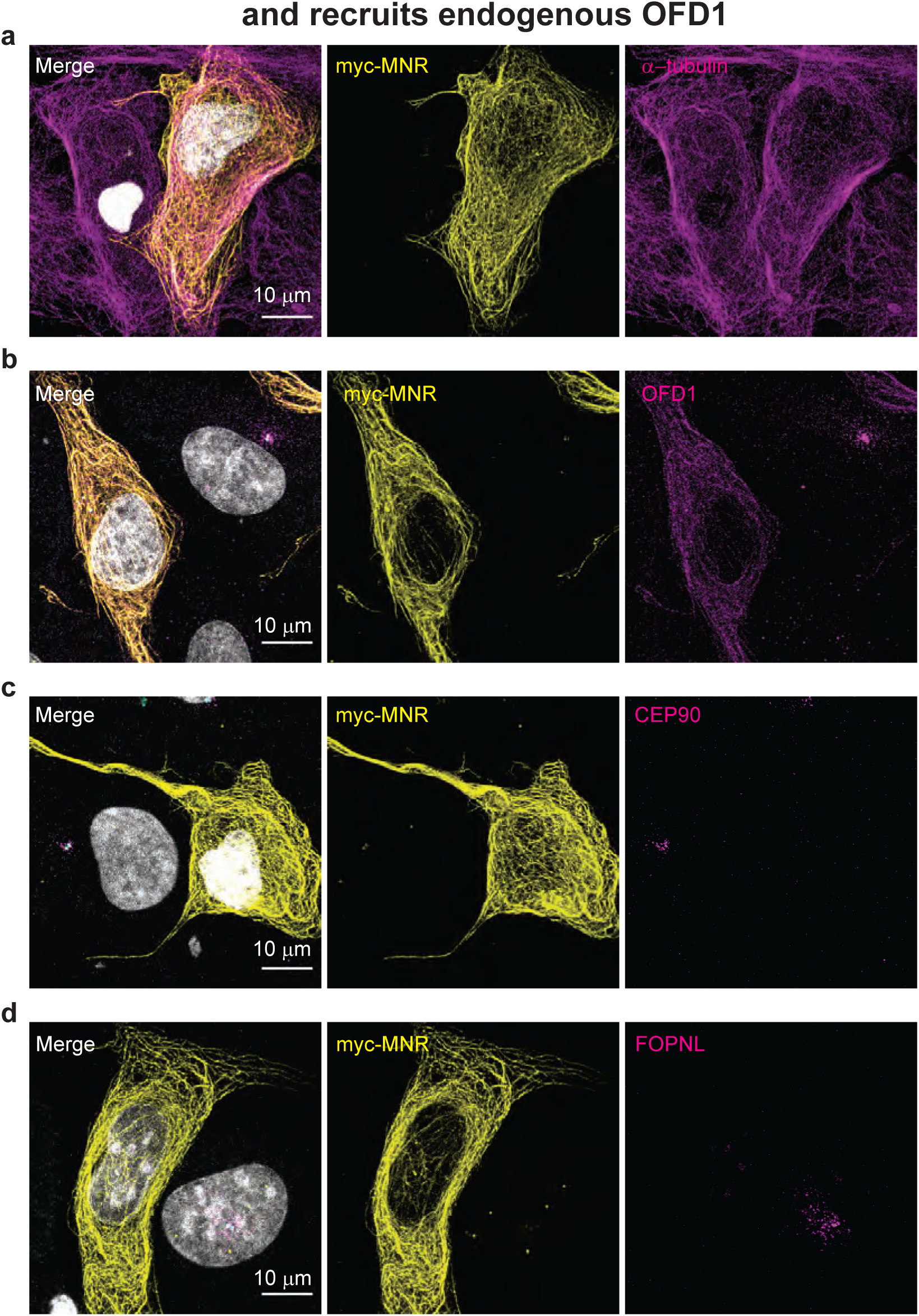
Overexpressed MNR localizes to microtubules and recruits endogenous OFD1. **a.** RPE1 cells transiently transfected with myc-tagged MNR and immunostained with antibodies to myc-tag and α-tubulin. Scale bar = 10 μm. **b.** RPE1 cells transiently transfected with myc-tagged MNR, and immunostained with antibodies to myc-tag and OFD1. Scale bar = 10 μm. **c**. RPE1 cells transiently transfected with myc-tagged MNR, and immunostained with antibodies to myc-tag and CEP90. Scale bar = 10 μm. **d**. RPE1 cells transiently transfected with myc-tagged MNR, and immunostained with antibodies to myc-tag and FOPNL. Scale bar = 10 μm.

**SUPPLEMENTARY FIGURE 3.**
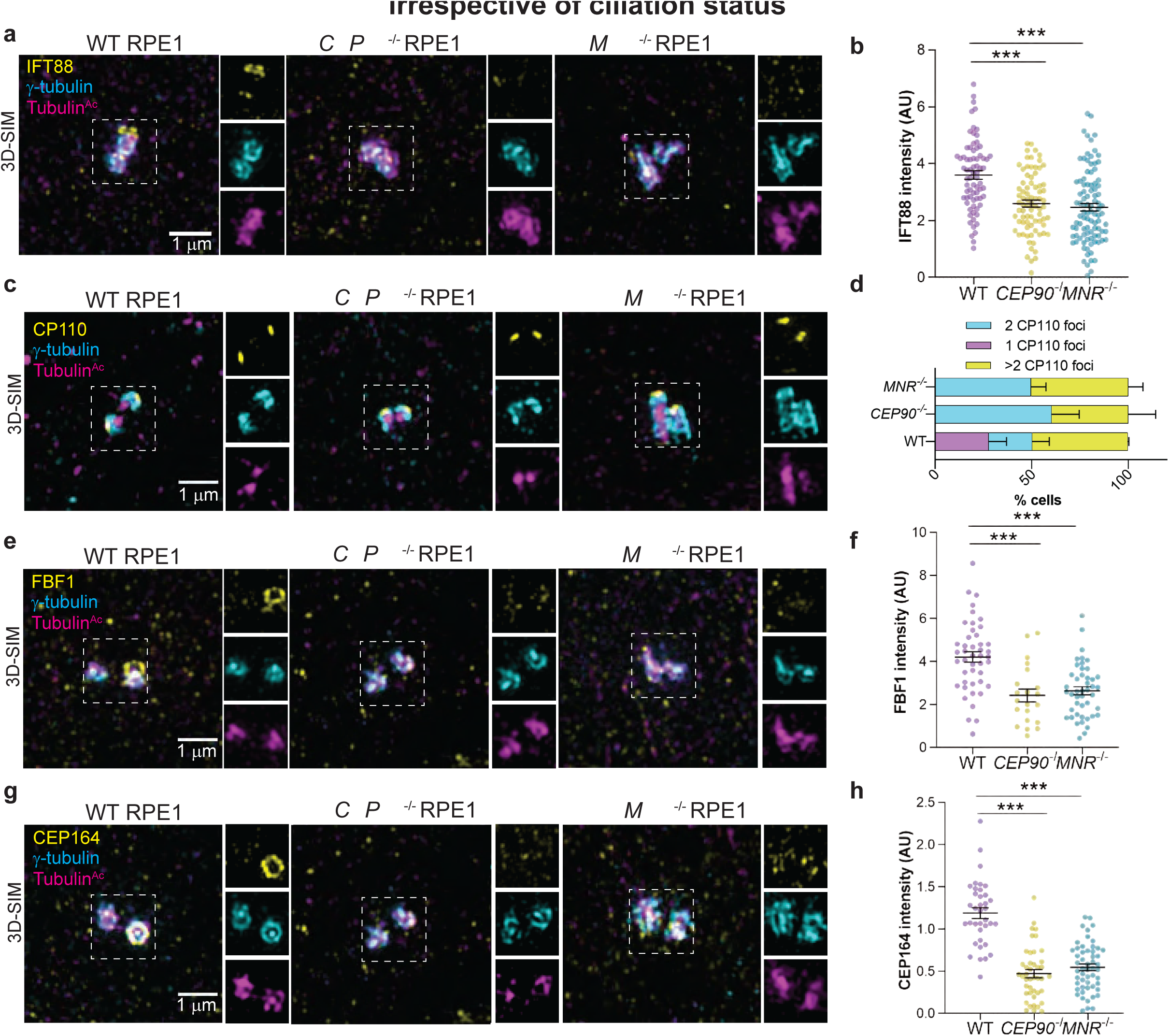
CEP90 and MNR regulate distal appendage assembly irrespective of ciliation status. **a**. Cycling WT, *CEP90*^-/-^ and *MNR*^-/-^ RPE1 cells stained with antibodies to IFT88, γ-tubulin (centrosome marker) and Tubulin^Ac^ (cilia and centriole marker). 3D-SIM imaging reveals IFT88 at the centrosome in WT cells, but not in *CEP90*^-/-^ and *MNR*^-/-^ cells. Scale bar for main panel and insets = 1 μm. **b**. Quantification of IFT88 fluorescence intensity at centrioles. Scatter dot plots show mean ± SEM. Asterisks indicate p<0.001, determined using one-way ANOVA. n = 73-96 measurements. **c**. Cycling WT, *CEP90*^-/-^ and *MNR*^-/-^ RPE1 cells immunostained for CP110 (yellow), centrioles (FOP, cyan) and Tubulin^Ac^ (cilia and centriole marker, magenta). Scale bar = 1 μm. **d**. Quantification of CP110 foci. **e**. Cycling WT, *CEP90*^-/-^ and *MNR*^-/-^ RPE1 cells stained with antibodies to FBF1, γ-tubulin (centrosome marker) and Tubulin^Ac^ (cilia and centriole marker). 3D-SIM imaging reveals FBF1 at the mother centriole in WT cells, but not in *CEP90*^-/-^ and *MNR*^-/-^ cells. Scale bar for main panel and insets = 1 μm. **f**. Quantification of FBF1 fluorescence intensity at centrioles. Scatter dot plots show mean ± SEM. Asterisks indicate p<0.001, determined using one-way ANOVA. n = 21-45 measurements. **g**. Cycling WT, *CEP90*^-/-^ and *MNR*^-/-^ RPE1 cells stained with antibodies to CEP164, γ-tubulin (centrosome marker) and Tubulin^Ac^ (cilia and centriole marker). 3D-SIM imaging reveals CEP164 at one of the two centrioles in WT cells, but not in *CEP90*^-/-^ and *MNR*^-/-^ cells. Scale bar for main panel and insets = 1 μm. **h**. Quantification of CEP164 fluorescence intensity at centrioles. Scatter dot plots show mean ± SEM. Asterisks indicate p<0.001, determined using one-way ANOVA. n = 37-53 measurements.

**SUPPLEMENTARY FIGURE 4.**
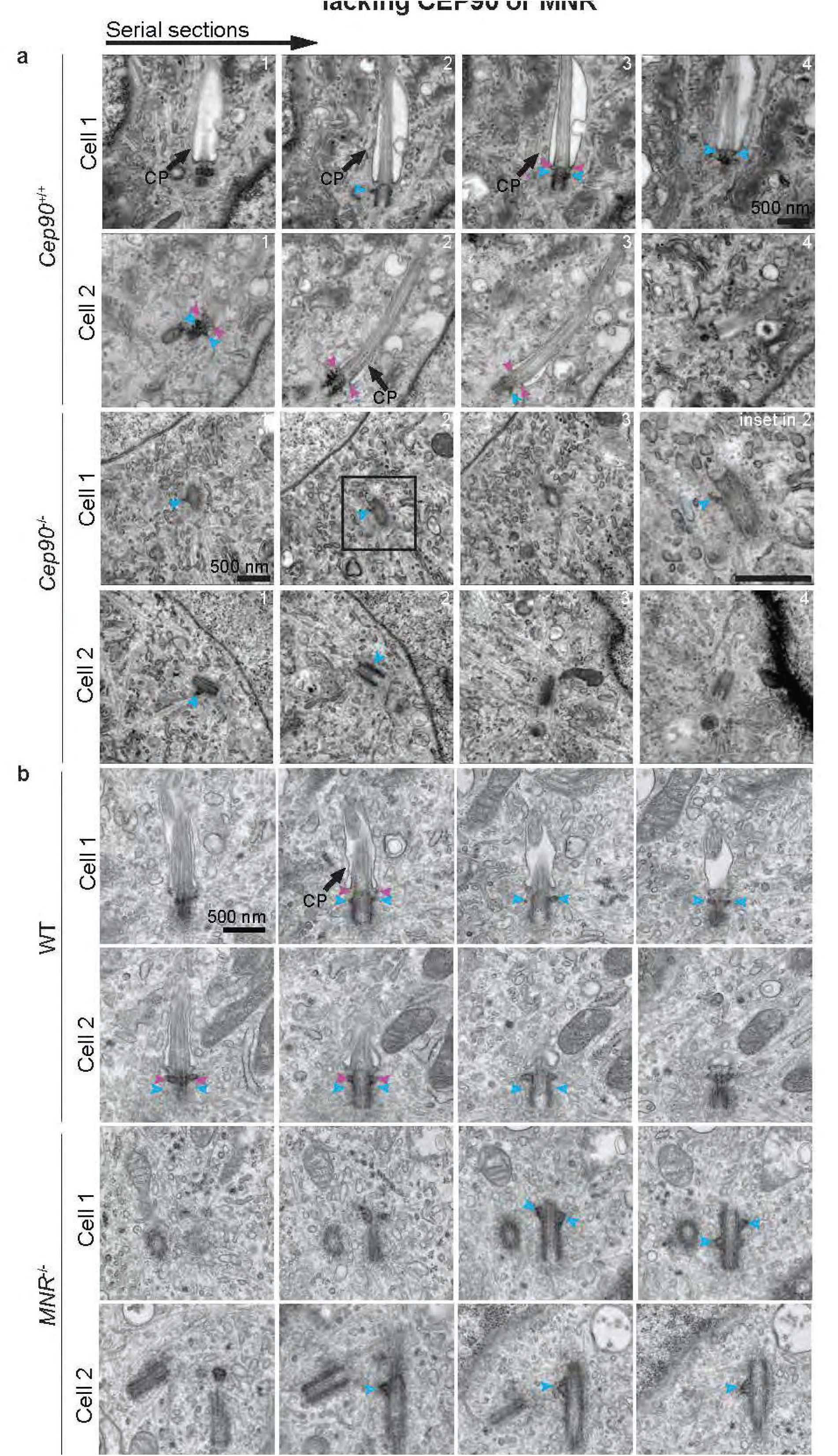
Serial section TEM confirms loss of distal appendages in cells lacking CEP90 or MNR. **a.** TEM images of*Cep90*^+/+^ and *Cep90*^-/-^ MEF cells confirms the absence of preciliary vesicle vesicle docking, and distal appendages at the *Cep90*^-/-^ mother centriole. Scale bar = 500 nm. Blue arrowheads indicate subdistal appendages, and pink arrowheads indicate distal appendages. CP indicate ciliary pocket. **b.** TEM images of WT and *MNR*^-/-^ RPE1 cells confirms the absence of preciliary vesicle docking and distal appendages at the *MNR*^-/-^ mother centriole. Centrioles are hyper-elongated in the absence of MNR. Scale bar = 500 nm. Blue arrowheads indicate subdistal appendages, and pink arrowheads indicate distal appendages. CP indicate ciliary pocket.

**SUPPLEMENTARY FIGURE 5.**
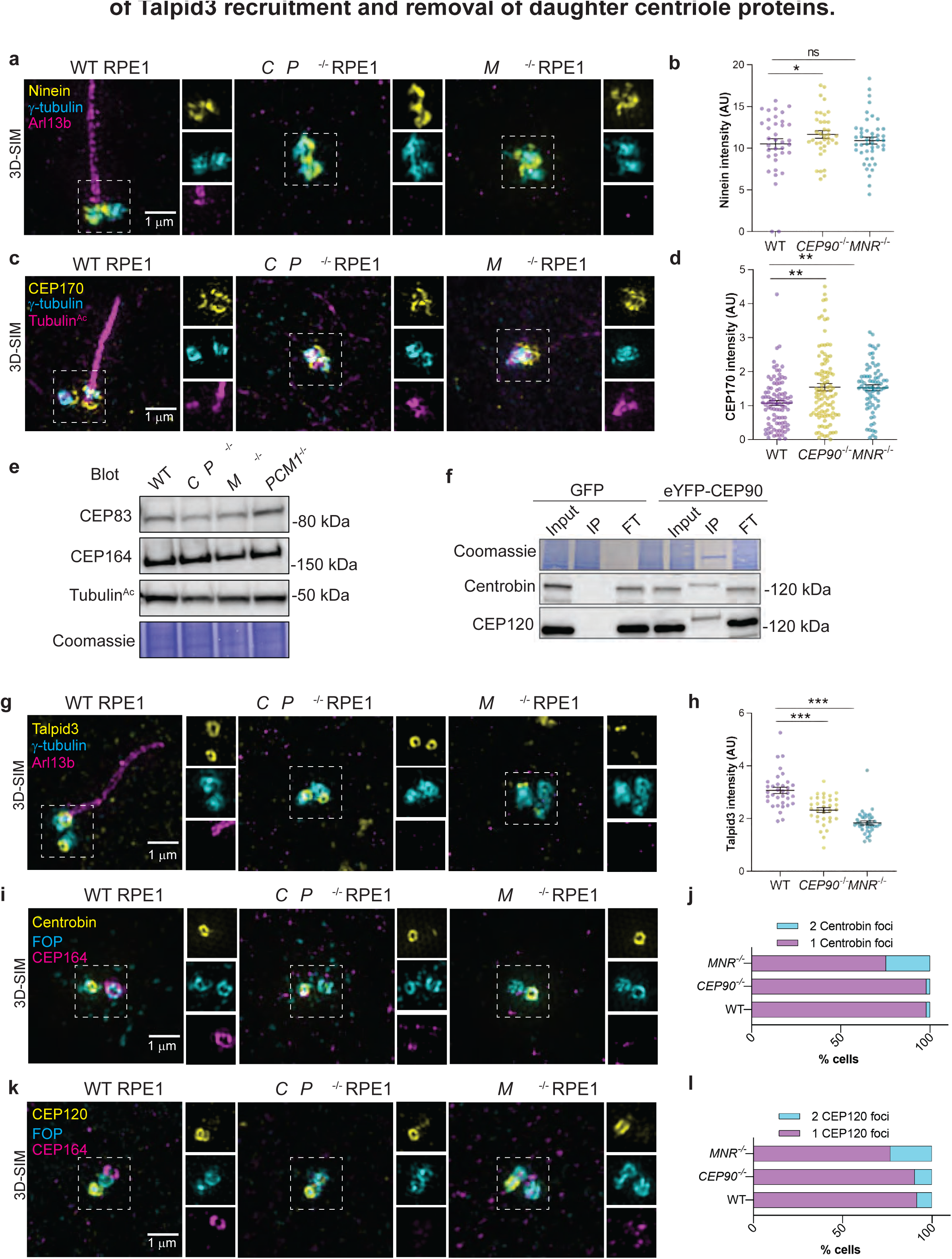
CEP90 regulates distal appendage assembly independent of Talpid3 recruitment and removal of Centrobin. **a.** 3D-SIM imaging of serum-starved WT, *CEP90*^-/-^ and *MNR*^-/-^ RPE1 cells immunostained for Ninein (yellow), centrioles (γ-tubulin, cyan) and cilia (ARL13B, magenta). Boxed regions are depicted in insets throughout. Ninein localizes to centrioles in WT, *CEP90*^-/-^ and *MNR*^-/-^ cells. Scale bar = 1 μm. **b**. Quantification of Ninein fluorescence intensity at centrioles in WT, *CEP90*^-/-^ and *MNR*^-/-^ cells. Horizontal lines indicate means ± SEM. Asterisks indicate p<0.05 determined using one-way ANOVA, ns indicates not significant. n = 36-45 measurements. **c**. 3D-SIM imaging of serum-starved WT, *CEP90*^-/-^ and *MNR*^-/-^ RPE1 cells immunostained for CEP170 (yellow), centrioles (γ-tubulin, cyan) and cilia (Tubulin^Ac^, magenta). Boxed regions are depicted in insets throughout. CEP170 localizes to centrioles in WT, *CEP90*^-/-^ and *MNR*^-/-^ cells. Scale bar = 1 μm. **d**. Quantification of CEP170 fluorescence intensity at centrioles in WT, *CEP90*^-/-^ and *MNR*^-/-^ cells. Horizontal lines indicate means ± SEM. Asterisks indicate p<0.05 determined using one-way ANOVA. n = 73-99 measurements. **e.** Immunoblot of distal appendage proteins CEP83 and CEP164 in WT, *CEP90*^-/-^, *MNR*^-/-^ and *PCM1*^-/-^ RPE1 cells. **f**. Co-immunoprecipitation of daughter centriole proteins Centrobin and CEP120 with CEP90. IP indicates eluate and FT indicates flow through **g**. WT, *CEP90*^-/-^ and *MNR*^-/-^ RPE1 cells were serum starved for 24 h and stained with antibodies to Talpid3, γ-tubulin (centrosome marker) and ARL13B (cilia marker). 3D-SIM imaging reveals ring of Talpid3 at mother and daughter centrioles. Scale bar = 1 μm for main panels and insets. **h**. Quantification of Talpid3 fluorescence intensity at centrioles. Scatter dot plots show mean ± SEM. Asterisks indicate p<0.001, determined using one-way ANOVA. n = 33-35 measurements. **i**. WT, *CEP90*^-/-^ and *MNR*^-/-^ serum-starved RPE1 cells immunostained for Centrobin (yellow), centrioles (FOP, cyan) and distal appendages (CEP164, magenta). Scale bar = 1 μm. **j**. Quantification of whether Centrobin localizes to one or two centrioles. n>50 cells from two biological replicates. CEP90 and MNR are not required to remove Centrobin from the distal mother centriole. **k**. WT, *CEP90*^-/-^ and *MNR*^-/-^ serum-starved RPE1 cells immunostained for CEP120 (yellow), centrioles (FOP, cyan) and distal appendages (CEP164, magenta). Scale bar = 1 μm. **j**. Quantification of whether CEP120 localizes to one or two centrioles. n>50 cells from two biological replicates. CEP90 and MNR are not required to remove Centrobin or CEP120 from the distal mother centriole.

---

**Table 1.** List of primary antibodies used in this study for immunofluorescence (IF) and western blotting (WB)

| Antibody | Host | Source | Catalog Number | Dilution/Fixation (IF) | Dilution (WB) |
| --- | --- | --- | --- | --- | --- |
| CEP90 | Rabbit | Proteintech | 14413-1-AP |  | 1:500 |
| CEP90 | Rabbit | Kunsoo Rhee<sup>1</sup> |  | 1:500/MeOH |  |
| MNR | Rat | Olivier Rosnet<sup>2</sup> |  | 1:500/MeOH |  |
| MNR | Rabbit | Novus Biologicals | NBP1-90929 |  | 1:500 |
| OFD1 | Rabbit | Novus Biologicals | NBP1-89355 | 1:500/MeOH | 1:500 |
| FOPNL | Rabbit | Sigma Prestige | HPA040599 | 1:100/MeOH |  |
| CEP164 | Goat | Santa Cruz Biotechnology | sc-240226 | 1:100/MeOH |  |
| CEP164 | Mouse | Novus Biologicals | NBP2-43707 |  | 1:2000 |
| γ-tubulin | Mouse | Sigma GTU-88 | T6557 | 1:500/MeOH | |
| Tubulin<sup>Ac</sup> | Rabbit | Cell Signaling Technology | 5335T | 1:2000/MeOH |  |
| Tubulin<sup>Ac</sup> | Mouse | Santa Cruz Biotechnology | sc-23950 AF647 | 1:100/MeOH |  |
| α-tubulin | Mouse | Sigma Aldrich | T5168 | 1:1000/PFA | |
| β-actin | Mouse | Proteintech | 6008-1-Ig | | 1:100,000 |
| GAPDH | Mouse | Proteintech | 60004-1-Ig |  | 1:100,000 |
| FBF1 | Rabbit | Proteintech | 11531-1-AP | 1:100/MeOH |  |
| ANKRD26 | Rabbit | GeneTex | GTX128255 | 1:100/MeOH |  |
| CEP83 | Rabbit | Sigma Prestige | HPA038161 | 1:100/MeOH |  |
| CENPF | Mouse | BD Biosciences | 610768 | 1:100/MeOH |  |
| Myo-Va | Rabbit | Novus Biologicals | NBP1-92156 | 1:100/MeOH |  |
| CP110 | Mouse | Millipore | MABT1354 | 1:100/MeOH |  |
| CEP97 | Rabbit | Proteintech | 22050-1-AP | 1:100/MeOH |  |
| IFT88 | Rabbit | Proteintech | 13967-1-AP | 1:100/MeOH |  |
| FOP | Rabbit | Proteintech | 11343-1-AP | 1:500/MeOH/ PFA |  |
| PCM1 | Rabbit | Proteintech | 19856-1-AP |  | 1:500 |
| PCM1 | Mouse | Santacruz | sc-398365 | 1:100/MeOH |  |
| Talpid3 | Rabbit | Proteintech | 24421-1-AP | 1:100/MeOH |  |
| Centrobin | Mouse | Abcam | Ab70448 | 1:100/MeOH | 1:1000 |
| Ninein | Rabbit | Michel Bornens | L79 | 1:200/MeOH |  |
| ODF2 | Mouse | Novus Biologicals | H00004957-M01 | 1:100/MeOH |  |
| CEP170 | Mouse | ThermoFisher | 72-413-1 | 1:100/ MeOH |  |
| Myc-tag | Goat | Novus Biologicals | NB600-335 | 1:500/PFA |  |
| Arl13b | Rabbit | Proteintech | 177100-1-AP | 1:500/MeOH/ PFA |  |
| mNeonGreen | Mouse | Chromotek | 32F6 | 1:500/PFA |  |
| Talpid3 | Rabbit | Proteintech | 24421-1-AP | 1:100/MeOH |  |
| CEP162 | Rabbit | Sigma Prestige | HPA030170 | 1:100/MeOH |  |
| CEP131 | Guinea pig | Kodani et al.,2015 <sup>3</sup> |  | 1:100/PFA/ MeOH |  |
| CEP120 | Rabbit | Betleja et al., 2018<sup>4</sup> |  | 1:3000/MeOH |  |

